# Small molecule FICD inhibitors suppress endogenous and pathologic FICD-mediated protein AMPylation

**DOI:** 10.1101/2024.07.13.603377

**Authors:** Bhaskar K. Chatterjee, Maroof Alam, Arghya Chakravorty, Shannon M. Lacy, Jason Rech, Charles L. Brooks, Peter D. Arvan, Matthias C. Truttmann

## Abstract

The AMP transferase, FICD, is an emerging drug target finetuning stress signaling in the endoplasmic reticulum (ER). FICD is a bi-functional enzyme, catalyzing both AMP addition (AMPylation) and removal (deAMPylation) from the ER resident chaperone BiP/GRP78. Despite increasing evidence linking excessive BiP/GRP78 AMPylation to human diseases, small molecules to inhibit pathogenic FICD variants are lacking. Using an *in-vitro* high-throughput screen, we identify two small-molecule FICD inhibitors, C22 and C73. Both molecules significantly inhibit FICD-mediated BiP/GRP78 AMPylation in intact cells while only weakly inhibiting BiP/GRP78 deAMPylation. C22 and C73 also efficiently inhibit pathogenic FICD variants and improve proinsulin processing in β cells. Our study identifies and validates FICD inhibitors, highlighting a novel therapeutic avenue against pathologic protein AMPylation.

## INTRODUCTION

Protein AMPylation is a post-translational protein modification (PTM) that regulates protein function by the covalent attachment of an AMP moiety to accessible hydroxyl groups of Thr, Ser and Tyr sidechains^1^. This ATP-dependent process is catalyzed by a dedicated set of enzymes called AMPylases. AMPylases can be broadly classified into two groups: enzymes that possess a highly conserved fic-domain (Fic)^2,3^, catalyzing the transfer of AMP, and non-Fic enzymes, such as SelO and DrrA, that catalyze AMPylation through a Fic-independent mechanism^4–6^.

This study focuses on the development of small molecular inhibitors specific for the AMPylase FICD, which regulates the Endoplasmic Reticulum (ER) heat shock 70 protein chaperone Binding immunoglobulin Protein (BiP)^7–9^.

Human FICD, also referred to as Huntingtin yeast-interacting partner E (HYPE) localizes to the ER lumen and is N-glycosylated on Asn275^8^. Structurally, FICD consists of a single transmembrane domain (residues 24–44), two TPR domains TPR1 (residues 105–135) and TPR2 (residues 140–170), and the conserved, catalytic Fic domain (residues 215–432) joined to the TPR motifs by a short linker (residues 170–215) (**Supplementary Figure S1**). The TPR motifs dictate FICD’s target recruitment^7,10–13^. The Fic core comprises the conserved catalytic loop and the flap^14^. The Fic core harbors the highly conserved Fic motif H_363_F(I/V)DGNGRT(S/A)R, while the flap (residues 311–324) is involved in positioning of the target residues. FICD possesses an auto-inhibitory helix (α-inh) containing the inhibitory motif (T/S)V(A/G)IE_234_N^11,15^. FICD catalyzes AMP transfer to target hydroxyl group(s) via the conserved His363 in the Fic motif, which acts as a base to attack the phosphodiester bond of an ATP molecule, resulting in AMP transfer and the concomitant release of a pyrophosphate group (PP_i_). Unlike most enzymes, FICD is bi-functional and catalyzes AMPylation as well as the removal of AMP from modified proteins (deAMPylation) using a single catalytic site^7,13,16^. The switch between AMPylation and deAMPylation states involves changes in enzyme oligomerization/monomerization, and an exchange of metal ions coordinating FICD’s active site^16^. Cellular signals that facilitate this switch remain poorly characterized but may involve changes in ER calcium levels^17^.

FICD regulates the ER stress response via reversible BiP AMPylation^7,13,18^. Published work is consistent with the model that under unstressed conditions, FICD AMPylates and generates a pool of primed (AMPylated) yet chaperoning-impaired BiP. The emergence of ER stress, however, results in rapid BiP deAMPylation, concomitant with the induction of the unfolded protein response (UPR^ER^)^19,20^.

Two recent studies describe pathologic *ficd* mutations with clinical implications because of dysregulated ER proteostasis^21,22^. Homozygous *FICD*^R371S^ expression in human patients is linked to infancy-onset *diabetes mellitus* and neurodevelopmental impairments^21^, whereas homozygous FICD^R374H^ expression leads to progressive motor neuron degeneration and peripheral neuropathy^22^. Both mutations cause FICD to lose its deAMPylation activity while slightly increasing or retaining AMPylation activity. This results in excessive BiP AMPylation impairing UPR^ER^ signaling with the concomitant accumulation of misfolded and aggregated polypeptides.

In this study, we develop a fluorescence polarization-based high-throughput screen to discover inhibitors of FICD-mediated protein AMPylation. We employ this platform to screen 84,480 small molecules obtained from two separate small molecule libraries and identify a total of 81 putative FICD inhibitors. Using orthogonal *in vitro* and cell-based assays, we identify two compounds (C22 and C73) that significantly inhibit endogenous FICD-mediated BiP AMPylation while weakly inhibiting BiP deAMPylation. C22 and C73 stably bind to the dimer interface of endogenous FICD and prevent the dimeric deAMPylase competent FICD from adopting an AMPylase competent conformation. We show that both compounds are non-cytotoxic small molecules that do not trigger the UPR^ER^ and are effective against pathogenic FICD mutants *in-vitro*. Finally, we demonstrate that C22 improves proinsulin folding and secretion in pancreatic β cells by reducing basal BiP AMPylation. Our study establishes FICD as a druggable target and suggests that targeting FICD may benefit multiple protein misfolding diseases.

## MATERIAL AND METHODS

### Protein expression and purification

Human His_6_-tagged _45-457_FICD constructs (WT, E234G, R371S and R374H) and *Caenorhabditis elegans* His_6_-tagged _187-457_FIC-1 were cloned into pETDuet-1 plasmids. The plasmids were transformed and expressed in *E. coli* BL21 or BL21-DE3 cells (Stratagene), and grown in TB medium containing 50 μg/mL of kanamycin to an optical density 0.8.-1. Protein expression was induced by adding 0.4 mM IPTG for 16–20 hours at 18 °C. Thereafter, bacteria were collected by centrifugation and bacterial pellets were sonicated in lysis buffer (50 mM HEPES, 250 mM NaCl, 10 mM imidazole, 1x protease inhibitor cocktail, pH 8.0). Lysates were cleared by centrifugation at 15,000 ×g for 30 mins. Supernatants were poured over nickel resin pre-equilibrated with lysis buffer. Thereafter, the resin was washed with wash buffer (50 mM HEPES, 250 mM NaCl, 30 mM imidazole, pH 8). His-tagged proteins were eluted in elution buffer (50 mM HEPES, 250 mM NaCl, 350 mM imidazole, pH 8). Fractions containing FICD were verified for purity by Sodium Dodecyl Sulphate-Polyacrylamide Gel Electrophoresis (SDS-PAGE), pooled, and dialyzed in a dialysis buffer overnight (50 mM HEPES, 150 mM NaCl, pH 8.0).

Human His_6_-SUMO-tagged BiP was expressed and purified as described previously^23^. Briefly, pSMT-WT BIP, kindly gifted by Dr. Liu (Virginia Commonwealth University), was expressed in *E. coli* BL21 cells grown in TB medium containing 50 μg/mL of kanamycin to an optical density of 0.6. Protein expression was induced by adding 1 mM IPTG for 5-6 h at 30 °C. Thereafter, bacterial pellets were sonicated in lysis buffer and centrifuged at 15,000× g for 30 mins. Supernatants were poured over nickel resin pre-equilibrated with lysis buffer. Thereafter, the resin was washed with a wash buffer. His-tagged proteins were eluted with an elution buffer (50 mM HEPES pH 8.0, 250 mM NaCl, 250 mM imidazole). Fractions containing BiP were verified for purity by Sodium Dodecyl Sulphate-Polyacrylamide Gel Electrophoresis (SDS-PAGE), pooled, and dialyzed in a dialysis buffer (50 mM HEPES, 150 mM NaCl, pH 8.0).

Purified FICD and BiP protein concentrations were measured spectrophotometrically at 280 nm using the Lambert-Beer law^24^. Thereafter, FICD and BiP aliquots were flash-frozen in liquid nitrogen and stored at −80 °C in storage buffer (50 mM HEPES, 150 mM NaCl, 10% (v/v) glycerol, pH 8.0).

### Fluorescence Polarization Assay

Binding kinetics of _45-457_FICD^E234G^ to the fluorescent ATP analog N6-(6-Aminohexyl)-ATP-5-FAM (FL-ATP) was determined by incubating increasing concentrations of the enzyme (0.75 μM - 2.5 μM), which was dissolved in an AMPylation buffer (50 mM HEPES pH 7.5, 150 mM NaCl, 10 mM MgCl_2_, 1mM EDTA and 1mM DTT), with 250 nM FL-ATP (final concentration). A Multidrop nano 384-well reagent dispenser (ThermoFisher, USA) was used to add FICD, dissolved in AMPylation buffer, and FL-ATP (dissolved in ultrapure MilliQ water) to a single 384-well black-bottom, black-walled microplate. The total reaction volume was 20 μL. The plate was snap-centrifuged at 1000× g for 60s and initial fluorescence polarization measurements were recorded using a BMG Pherastar plate-reader fitted with 485/530 nm filters before it was incubated in the dark at 37°C. Thereafter, the plate was loaded onto the plate reader and assessed for fluorescence polarization every 15 mins from the beginning of the incubation until 120 mins had elapsed. Samples containing FL-ATP in AMPylation buffer were used for setting the desired fluorescence gain adjustment.

### High-Throughput screening setup

A Multidrop nano 384-well reagent dispenser was used to pipette 2 μM of FICD^E234G^ dissolved in AMPylation buffer into columns 1–22. 2 μM _45-457_WT FICD was similarly added to column 23 as negative controls. FICD^E234G^ enzyme was then incubated with compounds or 1% (v/v) DMSO (positive control) for 10-15 mins at room temperature (RT). The Pintool Sciclone ALHD 3000 (Perkin Elmer) equipment was used to transfer 200 nL of DMSO-dissolved compounds from 2 mM source plates into 384-well black, flat-bottom, black-walled microplates, to obtain a final concentration of 20 μM in a total volume of 20 μL. 5120 compounds were sourced from the repurposing library (FDA approved drugs for other indications) maintained by the Centre for Chemical Genomics (CCG) at the University of Michigan. The compounds were added to columns 3–22 of each plate. Lastly, the multidrop reagent dispenser was used to pipette 1 μL FL-ATP (final concentration of 250 nM) into the whole plate. Plates were then incubated for 60 mins at 37°C in the dark. Post incubation, plates were loaded onto a BioTek stacker and scanned using the BMG Pherastar plate-reader, in succession, to obtain fluorescence polarization values using 485/530 nm filters.

Z’ and S/B (signal/background) values were determined by fitting the data to Equations (1) and (2), respectively,

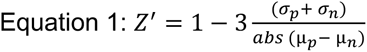

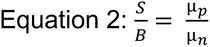

where µ_p_ and σ_p_ are the means and the standard deviations of the positive control samples and µ_n_ and σ_n_ are the means and the standard deviations of the negative control samples, respectively.

### Concentration-response curves (CRC)

A Multidrop nano 384-well reagent dispenser was used to pipette 1 μM FICD^E234G^ (positive control) or WT FICD dissolved in AMPylation buffer into designated microplate wells which already contained either 200nL of DMSO-dissolved compounds or an equivalent volume of 1% DMSO, respectively. Each compound was used at eight concentrations determined in accordance with semi-log fold dilutions starting from 30 nM. The reaction mixture was incubated for 10-15 mins at RT. 1 μL FL-ATP (final concentration of 250 nM) was dispensed using the automated reagent dispenser into the whole plate which was incubated for 90 mins at 37 °C in the dark. They were subsequently transferred to the BMG Pherastar plate-reader fitted with 485/530 nm filters to record fluorescence polarization. The same setup was used to obtain concentration-response curves for the commercially obtained Closantel analogs.

IC_50_ values were determined by fitting polarization values to Equation (3),

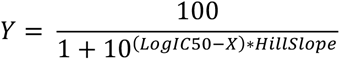

where Y is the polarization signal; X is the log concentration of inhibitor (μM) and IC_50_ is concentration of the inhibitor that elicits a response halfway between Bottom and Top. This is not the same as the response at Y=50. HillSlope describes the steepness of the family of curves.

### Tissue culture

A549 (ATCC-CCL 185) cells were grown in DMEM supplemented with 10% Fetal Bovine Serum (FBS) and 1% Penicillin-Streptomycin mixture (GM) at 37°C in 5% CO_2_ until they reached approximately 80% confluency. Cells were washed with PBS once, trypsinized, resuspended in GM, and plated in 6-well plates for assays.

Min6 (mouse insulinoma) cells were cultured in DMEM medium (25 mM glucose) supplemented with 10% FBS, 100 IU/mL penicillin and 100 μg/mL streptomycin, and 0.05 mM β-mercaptoethanol (GM6).

Neonatal primary cardiomyocytes were isolated from C57B6/J mice (post-natal day 1-3) as previously described^25^. Non-adherent cardiomyocytes were washed from the dish and re-plated in collagen pre-coated 96-well plates. Isolated cells were maintained in plating medium for 24 hours.

### BiP AMPylation kinetics in A549 cells

Cells were grown in 6-well plates. After the cells reached approximately 60% confluency, GM was removed, and cells were incubated for 15, 30, 45 and 60 mins in sterile PBS to assess BiP AMPylation levels.

When working with compounds, we preincubated cells with molecules C55, C83, C84, C522, C22, C73 and C34 in GM for approximately 12 hours. GM was removed and cells were exposed to PBS supplemented with the compounds for 60 mins. Cells preincubated with 0.5% (v/v) DMSO in GM or PBS served as negative controls.

When determining whether preincubation was sufficient for affecting BiP AMPylation, we preincubated cells with either 10 µM FICD inhibitors (C22 or C73) or DMSO, and subsequently exposed the cells to PBS for another 60 minutes supplemented with either the FICD inhibitors or DMSO only.

When determining whether supplementing FICD inhibitors in PBS is sufficient and necessary to affect BiP AMPylation, we did not preincubate the cells with compounds. Instead, cells were grown in GM until they reached approximately 70% confluency. Then, GM was removed, and PBS was added to the cells for 60 mins. FICD inhibitors or DMSO were added to PBS at 0 (immediately), 15 and 30 mins post PBS addition.

### BiP deAMPylation kinetics in A549 cells

Cells were grown to approximately 70% confluency. Next, cells were incubated with sterile PBS for 60 mins. Post incubation, PBS was removed, and GM was added for 1, 3, 5, 10 and 15 mins.

When working with FICD inhibitors, cells were incubated for 5 or 15 mins in GM supplemented with either 10 µM compounds (C22 or C73) or 0.5% (v/v) DMSO. Post GM incubation, cells were washed once with PBS to remove residual GM.

In all experiments, post incubation, cells were harvested and sonicated in cell-lysis buffer (20 mM Tris-HCl (pH 8.0), 100 mM NaCl, 1% NP-40, 2 mM EDTA and 1x protease inhibitor cocktail). The cell lysates were centrifuged at 10000x g for 10 mins at 4°C. The supernatant was carefully removed and used in a Bicinchoninic Protein Assay (BCA) to assess protein concentrations. 4× Laemmli sample loading buffer (BioRad) was added to the supernatant and the mixture was boiled for 5 mins at 95 °C. 10 µg of supernatant was loaded onto two separate 10% SDS-PAGE gels and resolved. Proteins were then transferred to PVDF membranes and blocked with 5% (w/v) milk or Bovine Serum Albumin (BSA) in Tris-buffered Saline supplemented with 0.1% (v/v) tween-20 (TBST). One of the membranes was blotted with mouse anti-Thr AMP (17G6, Biointron) while the other with anti-mouse BiP (Proteintech). Anti-mouse HRP-conjugated GAPDH (Proteintech) or mouse anti-α-Tubulin (Developmental Studies Hybridoma Bank) were used as loading controls. Membranes were incubated with primary antibodies (1:1000, diluted in TBST with 5% BSA or milk) at 4°C overnight and then incubated with HRP-conjugated-secondary antibody (1:5000) for 1 hour at RT. The membranes were incubated with ProSignal Dura ECL Reagent (Prometheus) at RT for 2 mins and imaged using Invitrogen iBright FL1500 Imaging system. Signals were quantified using ImageJ2 software^26^.

### Proinuslin secretion and folding

Min6 cells were grown to 80% confluency and then fed fresh GM6 supplemented with C22 (20 μM) or DMSO for 16 hours. Thereafter, both cells and media were collected, and cells were lysed in RIPA buffer supplemented with protease inhibitor cocktail. Cell lysates were clarified by centrifugation for 15 mins @ 12,000 rpm. Before electrophoresis, samples were boiled at (95 °C in SDS-gel sample buffer under either nonreducing or reducing (200 mM DTT) conditions, and then resolved either on straight nonreducing 15% SDS-PAGE in Tris-Glycine buffer or reducing 4-12% gradient NuPage gels. Proteins were then transferred to nitrocellulose membranes and blotted with mouse mAb anti-rodent proinsulin (Novus Biologicals), rabbit anti-BiP (Thermo) and mouse anti-AMPylated-BiP (17G6, Biointron). Mouse anti-β actin (Proteintech) was used as protein loading control.

Membranes were incubated with primary antibodies (1:1000, diluted in TBST with 5% BSA) at 4°C overnight and then incubated with HRP-conjugated-secondary antibody (1:5000) for 1 hour at RT.

### In vitro cytotoxicity assay

3-[4,5-dimethylthiazol-2-yl]-2,5 diphenyl tetrazolium bromide (MTT) assay was performed to determine the effect of compounds on the viability of HeLa (ATCC-CRM-CCL2), A549 (CCL-185), SK-N-SH (HTB-11) cells and neonatal murine cardiomyocytes. Immortalized cells were washed twice with PBS, trypsinized, resuspended in GM, and plated in a 96-well plate at a seeding density of 1.0 × 10^4^ cells/well. When the cells attained approximately 70% confluence, they were incubated with the indicated concentrations of C22 or C73 and kept at 37°C in 5% CO_2_ for approximately 24 hours. Cells incubated with 1% (v/v) DMSO alone served as negative controls. Final DMSO concentrations were kept below or at 1% (v/v). Post incubation, cells were washed once with sterile PBS. 10 μl of MTT solution (5 mg/ml in PBS) was mixed with 100 μl PBS, added to the cells, which were further incubated at 37°C in 5% CO_2_ for approximately 1-2 hours. The purple formazan crystals thus formed were dissolved in 100 μl of sterile DMSO, and the absorbance of the resulting mixture was measured using the Agilent BioTek Epoch 2 spectrophotometer at a sample wavelength of 540 nm and a reference wavelength of 630 nm. The Lethal Dose 50(LD_50_) values for C22 and C73 were determined by performing nonlinear regression analysis on the sigmoidal dose response curves obtained by fitting the data using GraphPad Prism (version 9.3.1, GraphPad Software). All experiments were carried out in triplicate.

### RNA isolation, processing, and quantitative PCR (qPCR)

Total RNA was isolated from 1 × 10^6^ A549 cells treated with 0.5 % (v/v) DMSO or 5 µM FICD inhibitors (C22 or C73) for approximately 24 hours using an RNA miniprep kit (Zymo Research) and quantified using a NanoDrop™ One Microvolume UV-Vis Spectrophotometer (Thermo Fisher). Cells exposed to 25 µg/ml tunicamycin, a well-known Endoplasmic Reticulum (ER) stress inducer, served as positive controls because tunicamycin induces the unfolded protein response (UPR) in the ER. 1 µg RNA was reverse-transcribed using the High-Capacity cDNA Reverse Transcription Kit (Applied Biosystems) to obtain complementary DNA (cDNA). 20 ng cDNA was mixed with ABclonal SYBR Green Master Mix (AbClonal) and primers for the human genes of interest (**Supplementary Table S1**), then plated in a single 384-well plate. Total volume of the reaction mixture was 20 µl. The plate was centrifuged at 600x g for 2 mins at RT and loaded onto a QuantStudio 5 Real-time PCR System. PCR conditions were determined following AbClonal protocol. Samples representative for each treatment, as described above, were pipetted in triplicate for each gene. Amplicons were quantified by comparison of 3-average ΔΔCT. Fold change in transcript levels were computed relative to *ACTB* respectively.

### In silico docking and molecular dynamics (MD) simulation

#### Rigid docking of C22 and C73 to FICD variants

The FASTDock program^27^ was used to identify putative binding sites on each of the target proteins discussed in our study. The default set of 18 chemical probes were used, and 2000 best docked poses were retained for each probe. For each probe, the top 5 clusters were considered for next steps. CDOCKER^28^ was used to dock compounds C22 and C73 at the top 5 putative binding sites. For each compound, the following docking protocol was applied: at the 5 binding sites, 10 different rotamers of each compound were generated using OpenBabel and docked while keeping the protein receptor region fixed and represented by a grid^28^. Then, the 10 poses were rescored by applying the Fast Analytical continuum Treatment of Solvation (FACTS) model^29^, which accounts for the desolvation penalty associated with each pose at the binding site. The FACTS-rescored docking scores were averaged across all poses. This protocol was independently repeated 10 times at each site for both compounds, thereby yielding an average dock score computed from a total of 100 poses for each compound. The site with the most favorable (most negative) averaged FACTS-rescored dock score was considered the top binding site, and the best binding pose at that site exhibited the highest (most negative) FACTS-rescored dock score. We generated 2D diagrams of protein-ligand interactions using PlexView (https://playmolecule.org/PlexView/).

### Molecular dynamics (MD) simulations

The stability of the FICD inhibitors at the previously obtained binding poses was probed using MD simulations. The CHARMM molecular simulation program^30^ was used to perform these simulations. Receptors with C22 or C73 bound were placed in a bath of explicit water models (TIP3P model) and counter ions (Na^+^ or Cl^-^) were added to neutralize the charge of the solvated system. The following scheme was used to perform all MD simulations performed in this study. First, the energy of the solvated system was minimized to eliminate bad contacts introduced during system preparation. During minimization, which included 2000 steps of steepest descent minimization followed by 1000 steps of ABNR minimization, the receptor and compound (ligand) heavy atoms were harmonically restrained to their initial poses with a force constant of 10 kcal/mol/ Å^2^. Next, an equilibration step was carried out under constant temperature (310 K) and volume conditions for 1 ns. The previous restraints were retained. Third, the restraints were reduced to 5 kcal/mol/ Å^2^ but were still imposed on the receptor and ligand heavy atoms, and the next phase of equilibration was carried out under constant temperature and constant pressure (1 atm) for another 1 ns. A penultimate phase of constant temperature and pressure (NPT) equilibration was initiated with restraint forces reduced to 1 kcal/mol/ Å^2^ for 1 ns. This was followed by another 1 ns of NPT equilibration with a very small restraint of 0.1 kcal/mol/ Å^2^. After these phases of equilibration, the production simulation was run for 5 ns under NPT conditions without any restraints.

The nonbonded interactions within the system were truncated at 12 Å after the application of potential switching function starting at 10 Å. Particle Mesh Ewald (PME) was used to treat long range electrostatics with *κ* = 0.32 Å^-1^, order = 4, grid size of 0.8-1.2 Å and force errors tolerance of 10^-5^. SHAKE^31^ was employed to constrain the distances of hydrogen-heavy atom bonds after performing hydrogen mass repartitioning^32^. The temperature was regulated using a Langevin thermostat with a friction coefficient of 5 ps^-1^. Pressure, in the constant pressure simulations, was isotopically regulated using a MC barostat with volume changes attempted every 25 steps. The integration timestep was set to 2 fs.

Force field parameters of the receptor(s), TIP3P water model and counter ions were taken from the CHARMM36 force field^33–35^. Parameters for the FICD inhibitors were obtained using the CGenFF program (v 2.5.1)^36^. The corresponding rtf files are shared as supplementary files S1 and S2.

### MM/PBSA calculations

Endpoint free energy calculations using the MM/PBSA technique were carried out using the snapshots generated from the production phase of the simulations. Only the last 4.5 ns of the 5 ns of the production was used to obtain snapshots every 40 ps. We used the single-trajectory protocol of MM/PBSA where the energy terms associated with the receptor-ligand complex, the receptor, and the ligand were all derived from the same snapshot as opposed to individual simulations conducted for each of these systems separately. The molecular mechanical (MM) energy terms and the surface area (SA) terms were determined using the CHARMM program. The SA term was used to compute the nonpolar component of the solvation energy by coupling it with a surface tension value of 0.005 kcal/mol/Å ^2^. The Poisson-

Boltzmann (PB) framework was used to compute the polar component. Delphi^37^ was used for the PB calculations. The internal and external dielectric values were set to 1 and 80, respectively. The salt concentration was set to 0 outside the solute’s (receptor, ligand or the complex) molecular surface, thereby reducing the Poisson-Boltzmann equation to Poisson equation only. The molecular surface itself was drawn using a solvent probe radius of 1.4 Å to emulate water. The solute atoms’ charges and radii were taken directly from the set used in the MD simulations.

### Statistical Analysis

Statistical Analysis was performed using the GraphPad Prism (version 10.2.2) software. Unpaired t-tests with Welch’s correction and 2-way ANOVA tests were performed. Figure legends specify the utilized tests for each data panel. p values were computed to determine statistical significance. If a p-value is less than 0.05, it is flagged with one star (*). If a p-value is less than 0.01, it is flagged with 2 stars (**). If a p-value is less than 0.001, it is flagged with three stars (***) and if a p-value is less than 0.0001, it is flagged with four stars (****).

All hit validation assays were performed at least in triplicate. For western blot-based quantifications, figures in the main manuscript show one representative western blot. The other replicates are shared in the supplementary figures.

## RESULTS

### An *in-vitro* fluorescence polarization screen identifies putative FICD inhibitors

To identify molecules that inhibit FICD-dependent protein AMPylation, we optimized an *in vitro* fluorescence polarization (FP) assay to measure FICD auto-AMPylation (**Supplementary Figure S2A**). In this assay, we used a fluorescent ATP analog N6-(6-Aminohexyl)-ATP-5-FAM (FL-ATP) (**Supplementary Figure S2B**) to measure auto-AMPylation of recombinant FICD^E234G^ (**Supplementary Figure S2C**). Unlike wild-type (WT) FICD, which purifies in an auto-inhibited conformation, FICD^E234G^ contains a dislodged α-inh domain, relieving auto-inhibition and turning this variant into a constitutive AMPylase^8,9,11^. To determine optimal reaction conditions, we tested different concentrations of FICD^E234G^ (**Supplementary Figure S2D**) and buffer components (**Supplementary Figure S2E**). We found that using 1 µM FICD and 0.25 µM FL-ATP in a buffer containing 50 mM HEPES pH 7.5, 150 mM NaCl, 10 mM MgCl_2_, 1mM EDTA and 1mM DTT provided a reliable dynamic window enabling to screen for both FICD^E234G^ activators and inhibitors. We next validated these reaction conditions comparing FP measurements of auto-inhibited FICD WT and FICD^E234G^ in a 384 well plate setup, which confirmed assay reproducibility and reliability (Z-factor = 0.83) (**Figure 1A**).

**Figure 1.**
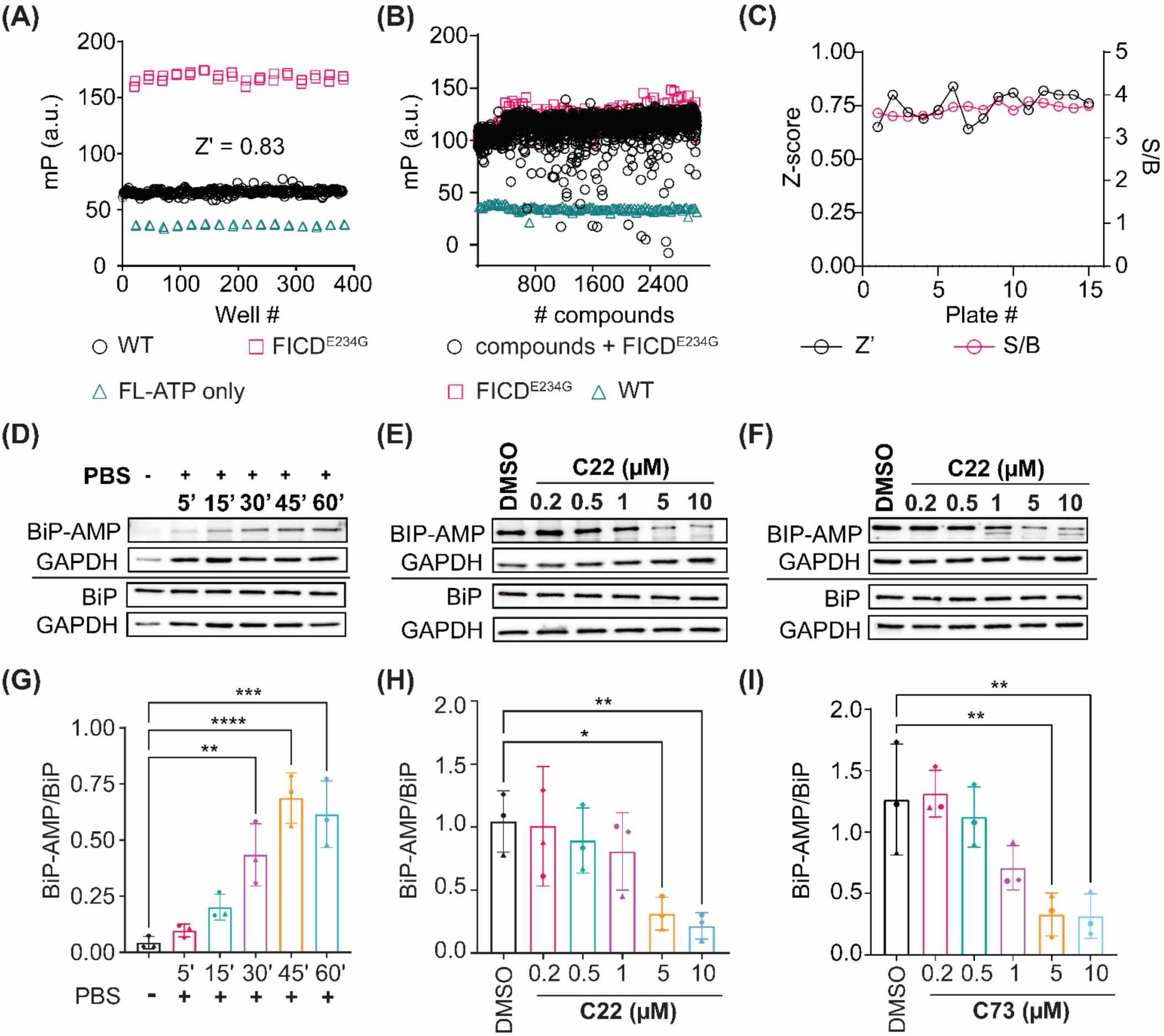
High-Throughput Screening (HTS) assay. (**A**) Assay variability was assessed by computing Z’. 2 µM WT or FICD^E234G^ diluted in AMPylation buffer was incubated with 250 nM FL-ATP. Wells containing only FL-ATP served as negative controls. (**B**) Randomized-well activity scatter plot of compounds from the repurposing library. Compounds were screened, in singlets (black), to identify putative FICD inhibitors. 1 µM WT FICD (red) or FICD^E234G^ (teal) was pipetted into a 384-well plate. Thereafter, compounds (at a final concentration of 20 µM) or DMSO (1% v/v) were added into test and control wells, respectively. Auto-AMPylation reaction was initiated by adding FL-ATP to each well and the reaction plate was incubated in the dark at 37°C for 90 mins. (**C**) Z’ scores and S/B ratio for 15 plates representing the repurposing library. (**D)** AMPylated and total BiP levels of A549 cells maintained in GM or treated with sterile PBS for the indicated time points. (**E**) Quantification of (**D**). **(F**) AMPylated and total BiP levels of A549 cells treated with PBS for one hour in the presence of either 0.5% (v/v) DMSO (control) or C22 (at indicated concentrations). (**G**) Quantification of (**F**). (**H**) AMPylated and total BiP levels of A549 cells treated with PBS for one hour in the presence of either 0.5% (v/v) DMSO (control) or C73 (at indicated concentrations). (**I**) Quantification of (**H**). Three independent biological replicas were analyzed for each timepoint for all cell-based assays. GAPDH was used as the protein loading control. A two-way ANOVA was performed to assess statistical significance between control and treated samples. Western blots of two remaining biological replicas for panels (**D**), (**F**) and (**H**) are shown in **Supplementary Figure S4.**

Using this optimized assay, we screened two distinct small molecule libraries: a commercially available (Selleck) repurposing library comprised of 5,120 FDA-approved compounds (**Figure 1B**), and a custom library consisting of 79,360 compounds from the Dart Neuroscience Collection. We maintained an average Z’ score of > 0.6 and an average signal-to-baseline ratio (S/B) of > 3 (**Figure 1C**) during our screen, offering a large dynamic window to identify potential FICD inhibitors. In our primary screen, we defined hits as compounds that inhibit FICD^E234G^ activity by >=20% or show a Standard Deviation (S.D.) >=3 from WT FICD activity as primary hits. Using these criteria, we identified 4,019 compounds in our primary screen (4.7% hit rate).

### Screening funnel confirms seven compounds as FICD inhibitors

To validate our primary hits, we followed a classic small molecule testing paradigm (**Supplementary Figure S3A**). First, we re-tested all 4,019 primary hits in a secondary screen, which was performed in triplicate.

These experiments confirmed 646 of the initial 4,019 compounds to reduce FICD^E234G^ auto-AMPylation. Of the 646 compounds, we selected 126 compounds that reduced FICD^E234G^ activity in both screens by at least 20%, showed good accessibility for site-chain modifications, and were not reported to exhibit pan-assay interference (PAINs). Of these 126 compounds, 55 belonged to the repurposing library and were tested in concentration-response experiments, in which 45 showed dose-dependent FICD inhibition (**Supplementary File 1**). The subsequent exclusion of known modulators of human enzymes further shrunk our hit list to 6 molecules.

Next, we used MScreen^38^, a compound management and HTS data analysis tool, to search for analogs exhibiting at least 40% structural similarity to the selected 6 hits. We identified 133 additional analogs, which were subsequently tested in triplicate. After these experiments, we were left with 19 compounds that reduced FICD^E234G^ auto-AMPylation by at least 20%.

Concentration-response experiments confirmed 13 of these 19 compounds as dose-dependent FICD^E234G^ inhibitors. We then repeated these tests using fresh commercially sourced chemical matter, which confirmed 7 compounds (**Supplementary Figure S3B**) exhibiting IC_50_ < 20 µM (**Supplementary Figure S3C, Supplementary Table 2**). These 7 molecules were selected for validation in orthogonal assays.

### Compounds C22 and C73 efficiently inhibit FICD-mediated BiP AMPylation in intact cells

Our initial *in-vitro* assays tested for inhibition of auto-AMPlyation of FICD^E234G^, a mutant enzyme version with constitutive AMPylation activity. In the next step, we determined if the seven putative FICD inhibitors reduced the activity of endogenous FICD in intact cells. In these assays, we analyzed AMPylation of BiP, perhaps the best characterized FICD target^8,9,18,19,22,39–42^. We incubated A549 cells with the compounds for approximately 12 hours in DMEM supplemented with 10% fetal bovine serum (GM). Afterwards, we replaced GM with sterile PBS containing the compounds and incubated the cells for another hour. In the absence of FICD inhibitors, the PBS treatment acutely depleted cells of nutrients, which led to a progressive increase in BiP AMPylation that reached saturation after approximately 45 mins (**Figure 1D and 1G**). In contrast, we observed that compounds C22 (**Figure 1E**) and C73 (**Figure 1F**) promoted a concentration-dependent inhibition of FICD-mediated BiP AMPylation in response to nutrition shortage. Compounds C47, C55, C83, C84, and C522 did not affect BiP AMPylation levels (**Supplementary Figure S4G**). At the highest tested compound concentration (10 µM), we observed up to 80% decrease in BiP AMPylation levels for both C22 (**Figure 1H**) and C73 (**Figure 1I**). These results confirm C22 and C73 as potent cell permeable FICD inhibitors.

### FICD inhibitors C22 and C73 have a favorable cytotoxicity profile and do not induce the UPR^ER^

Our cell-based assays showed that using compounds C22 and C73 at concentrations of 10 µM led to efficient FICD inhibition. To determine whether C22 or C73 are cytotoxic at micromolar doses, we incubated HeLa, A549, and SK-N-SH cells with increasing concentrations of each compound. We found that both compounds were well tolerated by all three immortalized cell lines, exhibiting LD_50_ (Lethal Dose_50_) values between 88 – 190.3 µM (**Supplementary Table 3**, **Supplementary Figure S5A-C**). Repeating the experiments using primary mouse cardiomyocytes, a cell type particularly sensitive to small-molecule toxicity^43–45^, (**Supplementary Figure S5D**) confirmed the limited cytotoxicity of compounds C22 and C73. Since FICD is a key regulator of the unfolded protein response in the endoplasmic reticulum (UPR^ER^)^7–9,46^, we next assessed whether C22 and C73 induce UPR^ER^ signaling. We performed quantitative PCR (qPCR) to evaluate expression levels of genes regulated by the UPR^ER^. We found that while the treatment with Tunicamycin, a known ER stressor^47,48^, induced a strong UPR^ER^, the exposure to C22 and C73 at inhibitory concentrations did not change the expression of selected genes involved in regulating ER homeostasis, inflammation, and apoptosis including *FICD*, *BiP*, Activating Transcription factor 4 (*ATF4*), spliced (s) and total (t) X-box Binding Protein 1 (*XBP1*), Activating Transcription Factor 6 alpha (*ATF6A*), *CASPASE4 and CASPASE6,* respectively^49^ (**Supplementary Figure S5E**). We also assessed CHOP protein levels in cells exposed to FICD inhibitors, which confirmed that neither FICD inhibitor promoted enhanced CHOP expression (**Supplementary Figure S5F**). These results provide strong evidence that FICD inhibitors C22 and C73 are tolerated across cell types at concentrations well outside the therapeutic window and do not disturb UPR^ER^-mediated cellular processes.

### C22 and C73 stably bind both monomeric and dimeric FICD *in-silico*

In unstressed cells, FICD is sought to reside preferentially in a deAMPylation-competent dimeric conformation. During our cell-based experiments, we exposed cells before and during nutrient depletion stress to FICD inhibitors. Thus, the compounds were likely interacting with both the deAMPylation-competent FICD dimer and, upon nutrient shortage, the AMPylation-competent FICD monomer. To understand whether FICD inhibitors show a preference for monomeric or dimeric FICD, we assessed their binding affinities to apo, dimeric WT FICD (PDB ID:4U04) and monomeric FICD^L258D^ (PDB ID:6I7J) by in-silico docking. First, we identify all potential binding sites in the WT dimeric FICD using FASTDOCK^50^. Then, we used CDOCKER^28^ to dock the FICD inhibitors to these binding cavities and rank the top 5 binding sites for both C22 and C73. (**Supplementary** Figure 6A). Two of these putative binding sites were particularly interesting: the smaller, potentially more flexible dimeric interface^11^ (site #1), and residues near the Tetratricopeptide repeat (TPR-II) domain (site #2) (**Supplementary** Figure 6B). C22 displays a stronger binding affinity (-28.86 +/- 4.89 kcal/mol) for site #1 than site #2 (-20.83 +/ 2.75 kcal/mol) while C73 displays similar binding affinities of -17.95 +/-5.25 kcal/mol and -15.232 +/9.99 kcal/mol to sites #1 and #2, respectively. Moreover, equilibrium MD simulations performed in the presence of explicit water molecules show that both compounds retain their respective binding modes with marginal deviations from the initial docked pose. This is evidenced by the fact that the structural Root Mean Square Deviation (RMSD) (of only the heavy atoms) for both compounds did not exceed 6.5 Å from the initial docked poses for majority of the MD simulation time-course (**Supplementary Figure 6C**).

Molecular Mechanics Poisson-Boltzmann Surface Area (MM/PBSA) calculations are considered more accurate in estimating binding free energies of protein-ligand complexes than scoring algorithms of most docking programs^51^. Hence, we used MM-PBSA scores as indicators of absolute binding energies for poses obtained from MD simulations of our docked FICD-inhibitor complexes. We found that residues near the TPR-II domain, which previously constituted the 2^nd^ ranked site for C22, displayed approximately two-fold higher MMPBSA score of –28.70 +/- 4.91 kcal/mol as compared to the smaller dimeric interface (–14.74 +/- 5.02 kcal/mol). This change can be attributed to a stronger binding mode because of the cation-π interactions between the guanidinium group on the sidechain of ARG180 (packed against the TPR-II domain) and the two aryl groups of C22 (**Figure 2A**). In contrast, the only prominent interaction is a hydrogen bond (H-bond) between the nitrile group in C22 and SER288 of one of the monomers at site #1 (**Figure 2A**), which could explain the lower affinity for that site. C73, the deprotonated form of C22, in accordance with our previous docking results, shows slightly favorable binding at site #2 (–269.61 +/- 16.70 kcal/mol) compared to site #1 (-256.38 +/- 7.31 kcal/mol). This slightly increased affinity is a result of the favorable cation-π interactions as well as a H-bond with the ARG180 sidechain at site #1 compared to H-bonds with the sidechains of ARG308 and ARG293 via its nitrile group at site #2 (**Figure 2B**).

**Figure 2:**
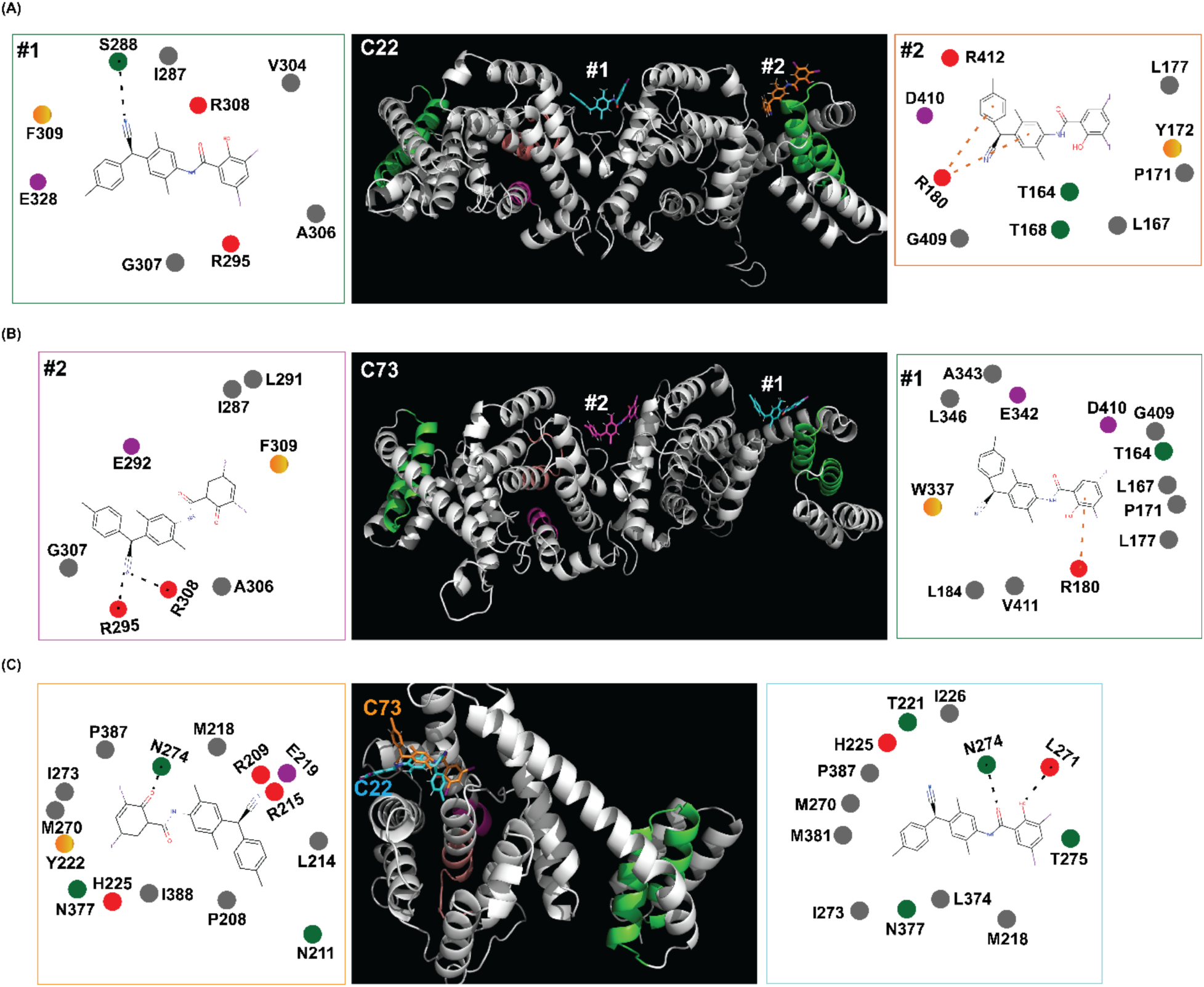
*In silico* docking and MD simulations of FICD inhibitors bound to dimeric and monomeric FICD. (**A-B**) The top two putative binding sites on the apo dimeric WT FICD (PDB ID: 4U04) for C22 (**A**) and C73 (**B**), respectively. # 1 and # 2 denote the top 2 sites. The inhibitory helix, catalytic core and TPR-II domains are highlighted in light magenta, salmon and green respectively. Prominent contacts between the compounds and their neighboring residues at these sites are shown in the cartoon representation. Contacts were drawn using PlexView and derived from the snapshot with maximum contacts across the last 100 snapshots (amounting to the first 4.5 ns of production simulation time). (**C**) The top binding site for C22 and C73 to the monomeric FICD^L258D^ (PDB ID: 6I7J). Prominent contacts are illustrated on the side. Yellow-orange, gray, purple, red and green circles represent aromatic, apolar, negatively charged, positively charged and polar interacting residues. Dotted black lines and dotted orange lines represent hydrogen bonds and cation-π interaction respectively.

In the next step, we used the same protocol to identify and rank putative binding sites of C22 and C73 on the strictly monomeric FICD variant^16^, FICD^L258D^ (**Figure 2C**). We found that C22 bound monomeric FICD with a heightened affinity to the top binding site as indicated by a higher MM/PBSA score (-43.33 +/- 6.52 kcal/mol) as compared to the top 2 binding sites on the dimeric FICD. At this site, the oxygen atoms of C22 formed H-bonds with ASN274 and LYS271 (**Figure 2C**). These H-bonds, in conjunction with the lack of cation-π interactions as observed in the dimeric form, explained the stronger binding affinity for the monomeric FICD. In contrast, C73 displayed a reduced binding affinity to monomeric FICD at the top binding site compared to the top 2 sites on the dimer (–124.28 +/- 9.23 kcal/mol) and engages in a single H-bond with ASN274 (**Figure 2C**).

These results indicate that FICD inhibitors, C22 and C73, may stably bind to both monomeric and dimeric FICD, and inhibit WT FICD-mediated BiP AMPylation by either preventing the switch to the AMPylation competent monomeric state or abrogating BiP binding to FICD.

### C22 and C73 inhibit both deAMPylation and AMPylation competent FICD states in intact cells

To validate and confirm our *in-silico* results, we preincubated A549 cells with 10 µM C22 and C73 for approximately 12 hours, allowing them to bind and stabilize deAMPylation-competent WT FICD. We utilized both regular DMEM and commercially available serum-free cell growth media (Opti-MEM) to exclude that the fetal calf serum we used in GM reduced bioavailability of the inhibitors^52^. Afterwards, we exposed these cells to PBS containing DMSO or compounds at 10 µM concentration. We found that preincubating cells with the FICD inhibitors in Opti-MEM, prior to nutrient depletion, was sufficient to prevent BiP AMPylation. In contrast, cells preincubated with C22 or C73 in GM show no reduction in AMPylated BiP. Cells continuously exposed to C22 or C73 in both media showed significant reductions in AMPylated BiP levels (**Figure 3A-B**). These results are consistent with our *in-silico* results and confirm that both C22 and C73 are capable of stabilizing the deAMPylation competent FICD conformation, thereby preventing its switch to an AMPylation competent state. To determine if C22 and C73 further inhibited the AMPylase-competent FICD confirmation, we devised an experiment in which we delayed the addition of FICD inhibitors until after the induction of nutrient depletion stress. This delay provided the cells with enough time to at least initiate the conformational switch required to transition from FICD-mediated deAMPylation to AMPylation. The addition of either C22 or C73 at onset of PBS-induced nutrient shortage significantly reduces BiP AMPylation. Delaying inhibitor addition by 15 mins attenuated the decline in BiP AMPylation, while a 30 min delay in C22 or C73 addition had no effect on BiP AMPylation levels (**Figure 3C-D**). Taken as a whole, these results support a model in which C22 and C73 are acting during the transition phase when the enzyme is adopting an AMPylation-competent conformation. This could occur through at least two mechanisms: First, C22 and C73 could stabilize FICD in a deAMPylation-competent conformation and prevent the transition. Second, the compounds could prevent AMPylation-competent FICD from engaging in BiP AMPylation.

**Figure 3:**
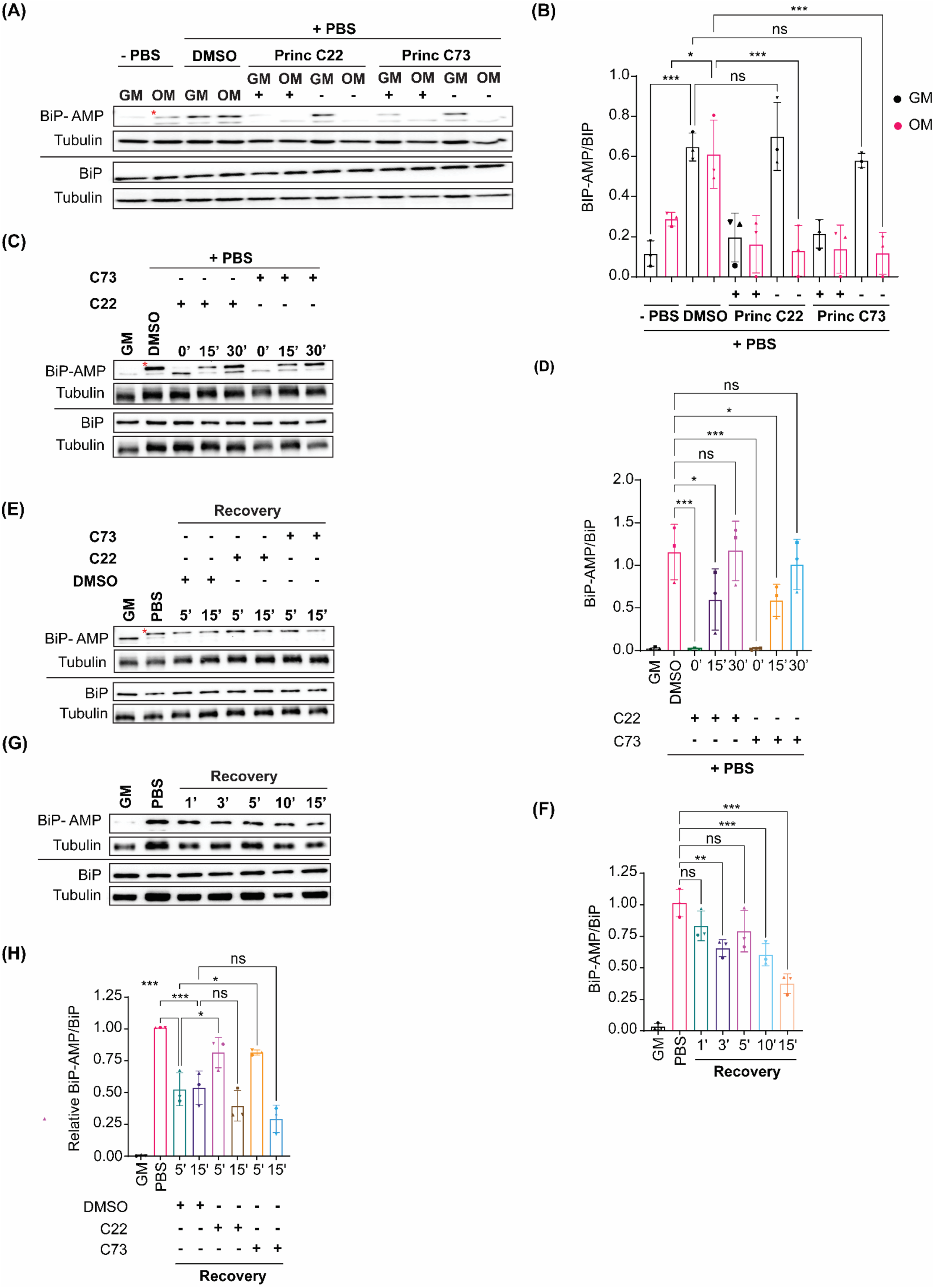
The effect of FICD inhibitors on deAMPylation and AMPylation competent FICD states. (**A**) AMPylated and total BiP levels in A549 cells preincubated with the 10 µM FICD inhibitors in GM or OM and subsequently exposed to PBS supplemented with 10 µM FICD inhibitors for 15 mins. Cells preincubated with 0.5% (v/v) DMSO and exposed to PBS supplemented with 0.5% (v/v) DMSO served as positive controls while cells grown in GM or OM supplemented with 0.5% (v/v) DMSO but not exposed to PBS served as negative controls. From left: Lanes 1-2 represent negative controls, lanes 3-4 represent positive controls, lanes 5-6 represent cells preincubated with C22 and exposed to PBS supplemented with C22 while lanes 7-8 represent cells preincubated with but not subsequently exposed to C22 during PBS treatment. Lanes 8-12 follow the exact order as lanes 5-8 but represent cells treated with C73. (**B**) Quantification of (**A**). (**C**) AMPylated and total BiP levels in A549 cells exposed to PBS supplemented with 10 µM C22 or C73 at the indicated time points and lysed 60 mins post PBS addition. Cells treated with 0.5% (v/v) DMSO in PBS for approximately 60 mins served as positive control. (**D**) Quantification of (**C**). (**E**) AMPylated and total BiP levels of PBS treated cells incubated in GM for the indicated time points. (**F**) Quantification of (**E**). (**G**) AMPylated and total BiP levels of PBS treated cells incubated in GM with or without 10 µM C22 or C73, for the indicated time points. (**H**) Quantification of (**G**). Three independent biological replicas were analyzed for each timepoint. α-Tubulin was used as the protein loading control. Cells treated with 0.5%(v/v) DMSO in PBS served as positive control. GM indicates cells never exposed to PBS which served as negative controls. BiP-AMP/BiP signal intensity ratios were computed relative to positive controls. A two-way ANOVA was performed to assess statistical significance between control and treated samples. * indicates the protein band used for quantification purposes; the lower protein band is non-specific for BiP AMPylation. Western blots of two remaining biological replicas for panels (**A**), (**C**), (**E**) and (**G**) are shown in **Supplementary Figure S7.**

### C22 and C73 only weakly inhibit BiP deAMPylation in intact cells

FICD is a bifunctional enzyme preferentially adopting a deAMPylase-compentent conformation in the absence of stress^19,20^. To further characterize the mode of action of compounds C22 and C73, we tested whether these molecules could inhibit FICD-mediated BiP deAMPylation in intact cells. Our assay was based on the finding that starvation mediated BiP AMPylation in A549 cells was quickly reversed upon incubating the cells in complete medium (**Figure 3E**). The incubation of nutrient depleted cells in GM for 5 minutes was sufficient to significantly reduce AMPylated BiP levels, with a maximal approximately 60% decrease occurring after 15 mins in complete medium (**Figure 3F**). The addition of both C22 and C73 delayed but did not prevent BiP deAMPylation from occurring (**Figure 3G-H**). These results indicate that C22 and C73 are weak modulators of FICD-mediated BiP deAMPylation in intact cells.

### C22 derivative C34 promotes BiP AMPylation inhibition potency in intact cells

C22, and its sodium salt, C73, are both halogenated salicylanilide molecules. To identify C22 analogs with improved FICD inhibition activity, we tested several commercially available analogues using our well-established FP assay (**Table 1**). Interestingly, the benzamide moiety was intolerant to most structural changes that we evaluated. The removal of the 3- and 5-iodo functional groups (compound 2 (C51)), resulted in an approximately 40-fold reduction in potency (**Table 1**, **Figure 8A**). The incorporation of a 5-chloro-2-hydroxy benzamide (compound 3 (C32)), modestly improved inhibition potency compared to C51 (**Table 1**, **Figure 4A**). Adding an unsubstituted benzamide (compound 4 (C50)), and 3-nitro-2-methyl benzamide, (compound 5 (C52)) resulted in complete loss of potency (**Table 1**, **Figure 4A**). While only a limited number of benzamide analogues were tested, compounds 2-5 indicate that the 2-hydroxyl moiety is essential for compound potency, and halogenated substitutions at the 3- and 5-positions are required for compounds to be effective FICD inhibitors. Of all six tested analogs, compound 6 (C34) was of particular interest. The replacement of racemic nitrile with a carbonyl moiety while retaining the 3- and 5-iodo groups resulted in C34 exhibiting significantly improved potency compared to compounds 2-5, and a similar potency compared to our benchmark compounds, C22 and C73 (**Table 1**, **Figure 4A**). In cell-based assays, we further observed that C34 promoted a concentration-dependent decrease in FICD mediated BiP AMPylation (**Figure 4B**). We observe approximately 75% decrease in BiP AMPylation levels at 1 µM and a near complete inhibition of BiP AMPylation at 5 and 10 µM (**Figure 4C**). This was superior to the FICD inhibition profile of both C22 (**Figure 1D**) and C73 (**Figure 1F)**, which reduced BiP AMPylation levels by approximately 80% at the highest tested concentration (10 µM). These results indicate functional group flexibility in the biaryl region of the scaffold could be further exploited to rationally improve these FICD inhibitors.

**Figure 4:**
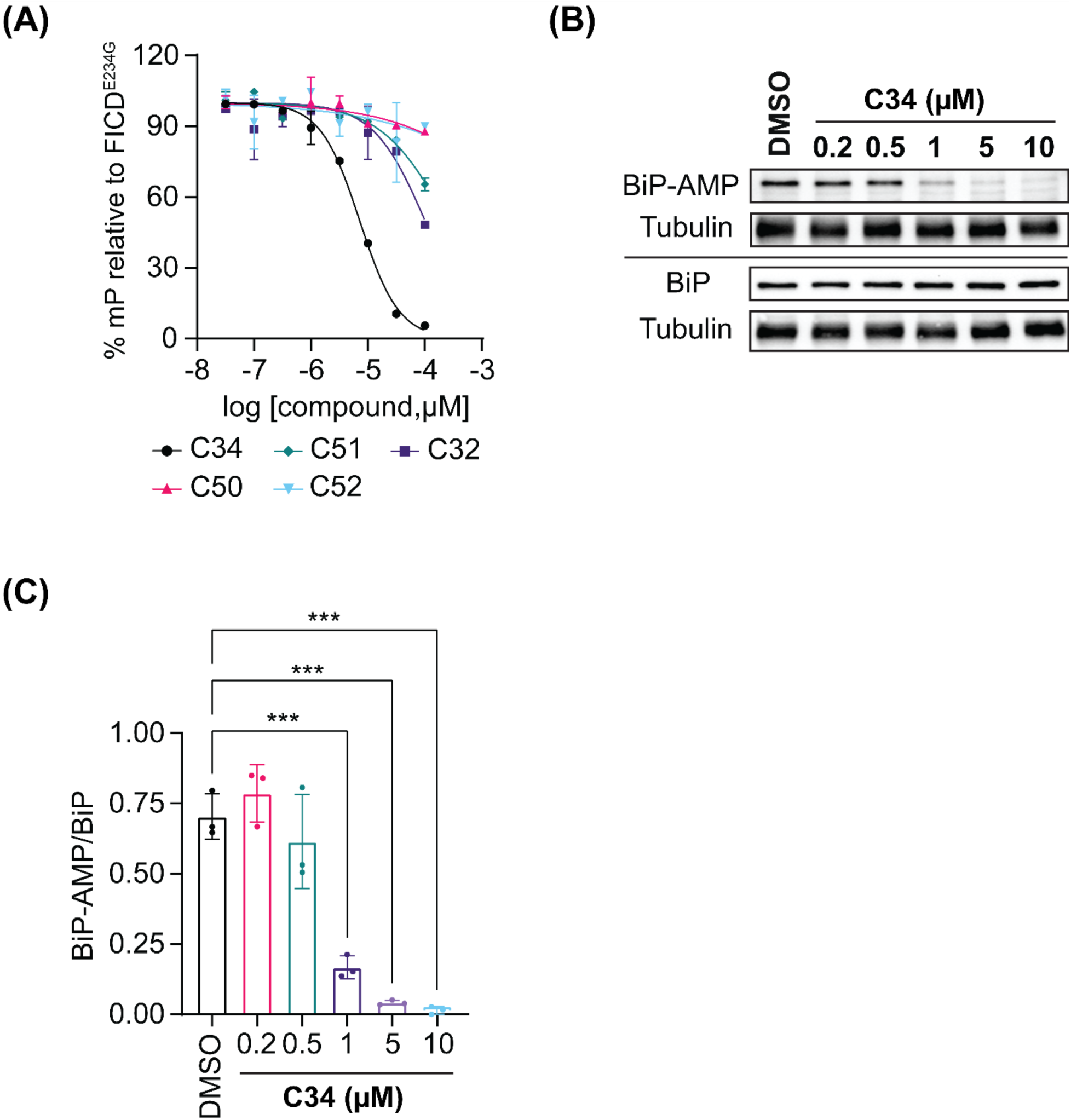
Concentration response assessment of compound C34. (**A**) 1 µM FICD^E234G^ was incubated with varying concentrations of C34 for 10-15 mins at RT, following which FL-ATP was added. The reaction was incubated for 90 mins at 37°C in the dark. Each dot represents the mean of duplicate mP measurements with the arrows representing standard error of mean (SEM). (**B**) AMPylated and total BiP levels of A549 cells treated with PBS for approximately 60 mins in the presence of either 0.5% (v/v) DMSO (control) or C34 (at indicated concentrations). Tubulin was used as the protein loading control. (**C**) Quantification of (**B**). Three independent biological replicas were analyzed for each timepoint. A two-way ANOVA was performed to assess statistical significance between control and treated samples. Western blots of two remaining biological replicas are shown in **Supplementary Figure S8.**

**Table 1:**
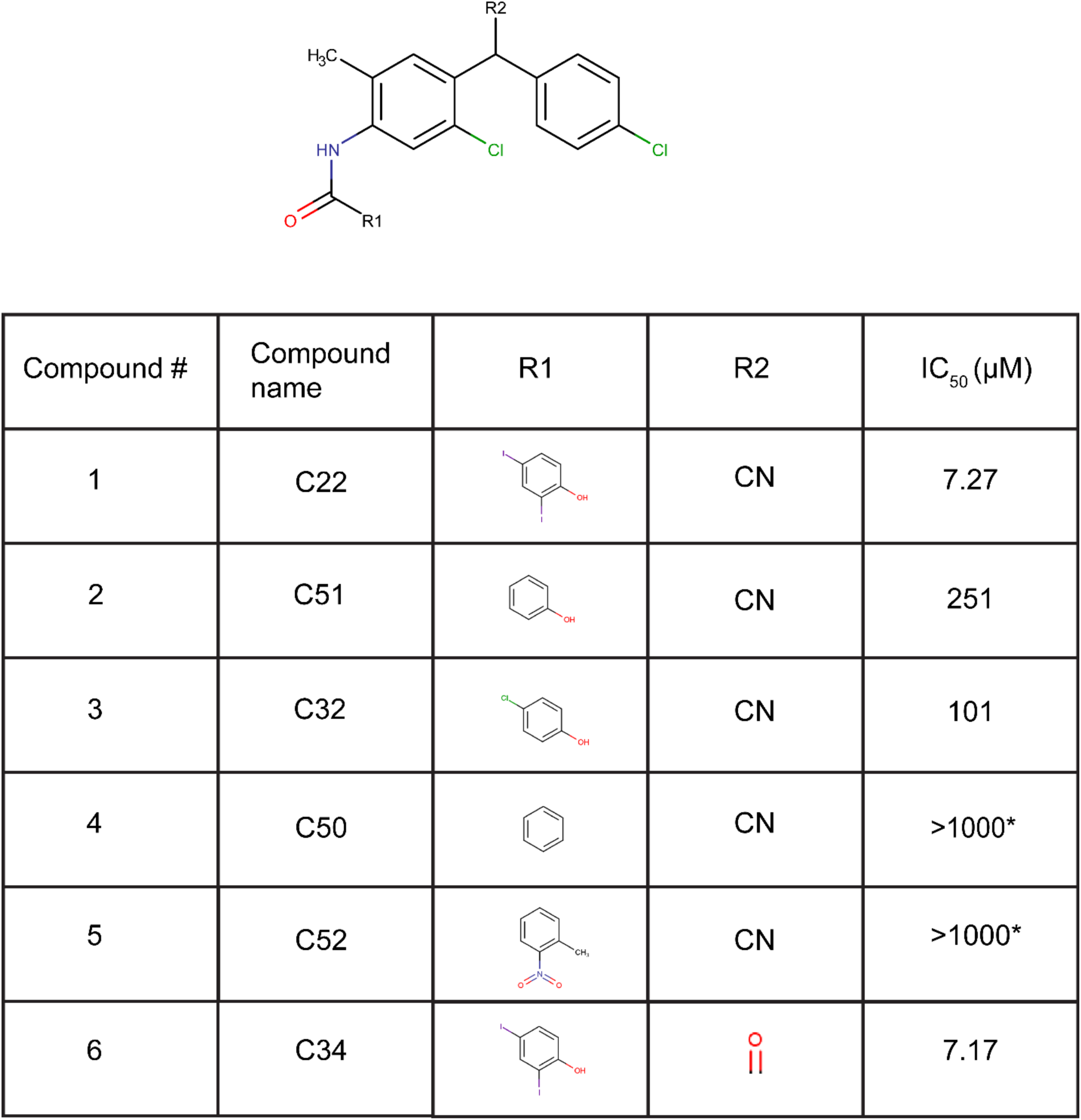
Commercially available closantel analogs. Compound 1 is closantel while compounds 2-6 are closantel derivatives. All compounds share the same core structure depicted above the table. IC50 values (µM), as shown in the table, were obtained by fitting the concentration response data using a non-linear regression method in GraphPad Prism. * indicates extrapolated values. The best fit curve should be interpreted with caution as it showed a low R-squared value.

### FICD inhibitors C22 and C73 inhibit human-pathogenic FICD variants *in vitro*

Recent reports link two mutations in the FICD active site to infancy-onset diabetes and motor neuron degeneration^21,22^. In both cases, the pathogenic FICD variants, FICD^R371S^ and FICD^R374H^, excessively AMPylate BiP. We thus tested whether FICD inhibitors C22 and C73 suppress FICD^R371S^ and FICD^R374H^ *in vitro*. Using our FP assay, we observed that both FICD^R371S^ (**Figure 5A**) and FICD^R374H^ (**Figure 5B**) showed auto-AMPylation activity, yet substantially less than FICD^E234G^. Incubating both pathologic FICD mutants with increasing concentrations of FICD inhibitors did not significantly decrease auto-AMPylation levels except at the highest tested concentration (100 µM) where C22 and C73 elicited auto-AMPylation reduction by approximately 30% and 20%, respectively (**Figure. 5C**). Despite limited inhibition of auto-AMPylation, both C22 and C73 significantly reduced BiP AMPylation *in vitro* at a molar ratio of 20:1 (inhibitor: protein) (**Figure 5D-E**). These results indicate that Arg371Ser and Arg374His mutations have a distinct impact on FICD auto- and BiP AMPylation and define C22 and C73 as potent inhibitors of pathologic FICD variants.

**Figure 5:**
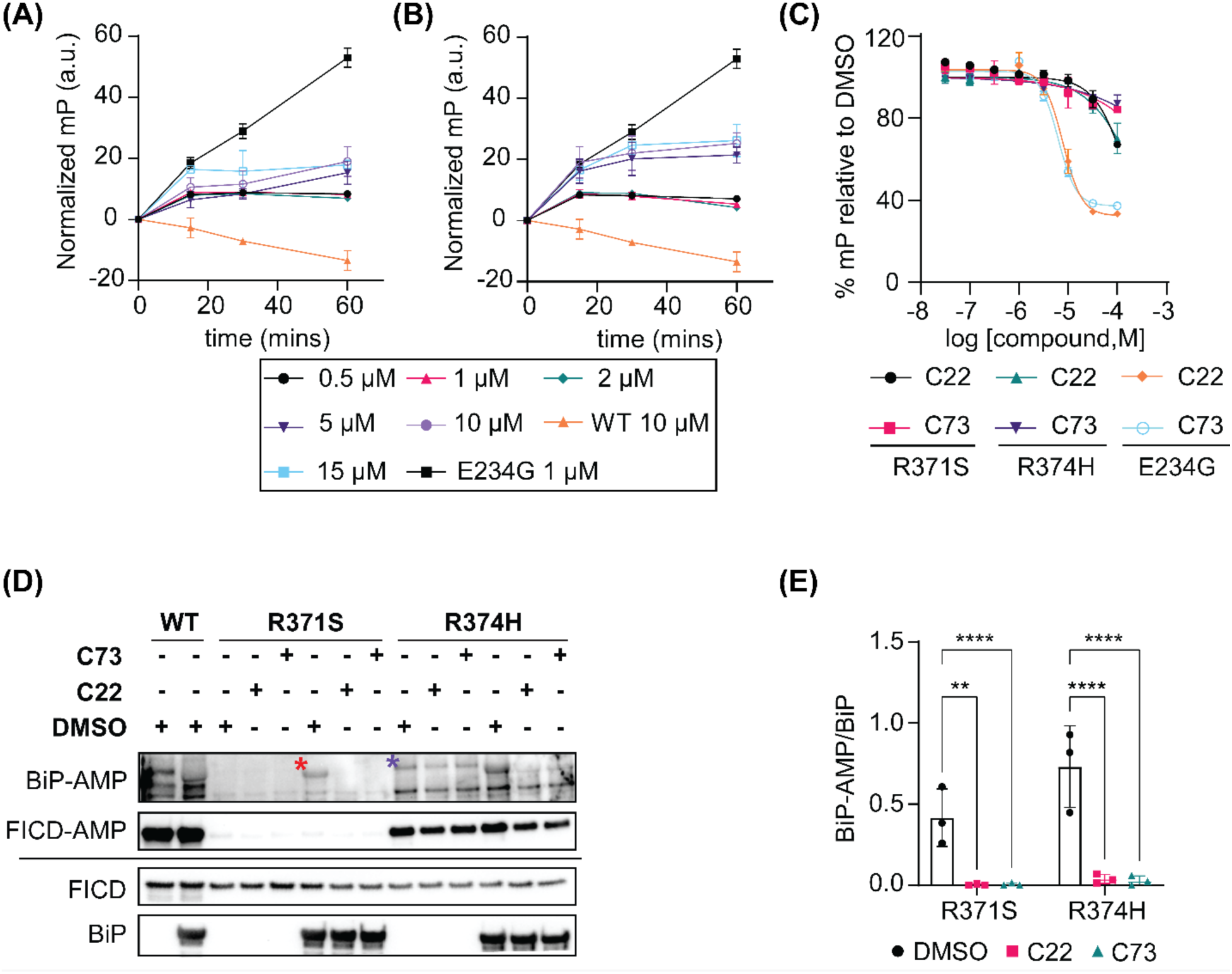
Effect of C22 and C73 on pathologic protein AMPylation *in-vitro*. Auto-AMPylation activity of (**A**) FICD^R371S^ and (**B**) FICD^R374H^ was assessed by incubating FICD variants at indicated concentrations with 250 nM FL-ATP. FP was measured at indicated timepoints. Wells containing WT FICD and FICD^E234G^ served as negative and positive controls respectively. Triplicate measurements were averaged for each FICD concentration and plotted as a function of time. (**C**) 1 µM FICD^E234G^ and 5 µM FICD^R371S^ and FICD^R374H^ were incubated with varying concentrations of C22 and C73 for one hour and FP measured thereafter. FICD mutants incubated with DMSO served as negative controls. Triplicate measurements were averaged for each compound concentration and plotted for each FICD mutant. (**D**) AMPylated and total levels of recombinant human BiP in the presence of FICD inhibitors or DMSO. WT FICD (+/-) BiP served as negative controls. * indicates the protein band used for quantification purposes while * indicates non-specific AMPylation of a protein contaminant which co-purified with recombinant FICD. (**E**) Quantification of (**D**). Three independent biological replicas were analyzed for each timepoint. A two-way ANOVA was performed to assess statistical significance between control and treated samples. Western blots of two remaining biological replicas are shown in **Supplementary Figure S9.**

### C22 promotes proinsulin folding and secretion in pancreatic β-cells

Recent work suggests a novel link between infancy-onset diabetes and excessive FICD-mediated BiP AMPylation^21^. Since BiP’s chaperone function is required for proinsulin folding^53^ and FICD-mediated BiP AMPylation inhibits its chaperone activity^8,9,13,46^, we investigated whether reducing BiP AMPylation using FICD inhibitors improves proinsulin folding and secretion in Min6 pancreatic β-cells were treated with 20 μM FICD inhibitors (inhibitor concentration was optimized as shown in **Supplementary Figure S10A-B)** or DMSO for 16 hours, and both conditioned media (M) and cell lysates (C) were probed for proinsulin levels (**Figure 6A**). Proinsulin protein content in the complete system (M+C) was unaffected by the 16 h treatment with both C22 (**Figure 6B**) and C73 (**Supplementary Figure S10C**). Interestingly, we observed an approximately 8-fold increase in proinsulin levels in M as compared to C when cells were treated with C22 (**Figure 6C**) and a roughly 2-fold increase when cells were exposed to C73 (**Supplementary Figure S10D**). To elucidate the reason for such an observation, we investigated whether C22 affected proinsulin protein folding by measuring the abundance of aberrant disulfide-linked proinsulin complexes. Two of the most readily quantifiable misfolded forms of proinsulin are its disulfide-linked dimer and trimer forms, and the ratio of these aberrant forms to monomeric proinsulin was markedly improved by C22 treatment (**Figure 6D-E**). We hypothesized that C22 mediated reduction in BiP AMPylation may contribute to improved proinsulin folding by increasing the pool of active BiP. Indeed, we observed an approximately 2.5-fold decrease in AMPylated BiP levels in cells treated with C22 as compared to cells treated with DMSO (**Figure 6F-G**) while cells treated with C73 showed no changes in AMPylated BiP levels (**Supplementary Figure S10E-F**). These results indicate that FICD inhibitor C22 promotes proinsulin folding and anterograde trafficking out of the ER by increasing the pool of chaperone-competent BiP, which reduces misfolded or aggregated proinsulin levels.

**Figure 6:**
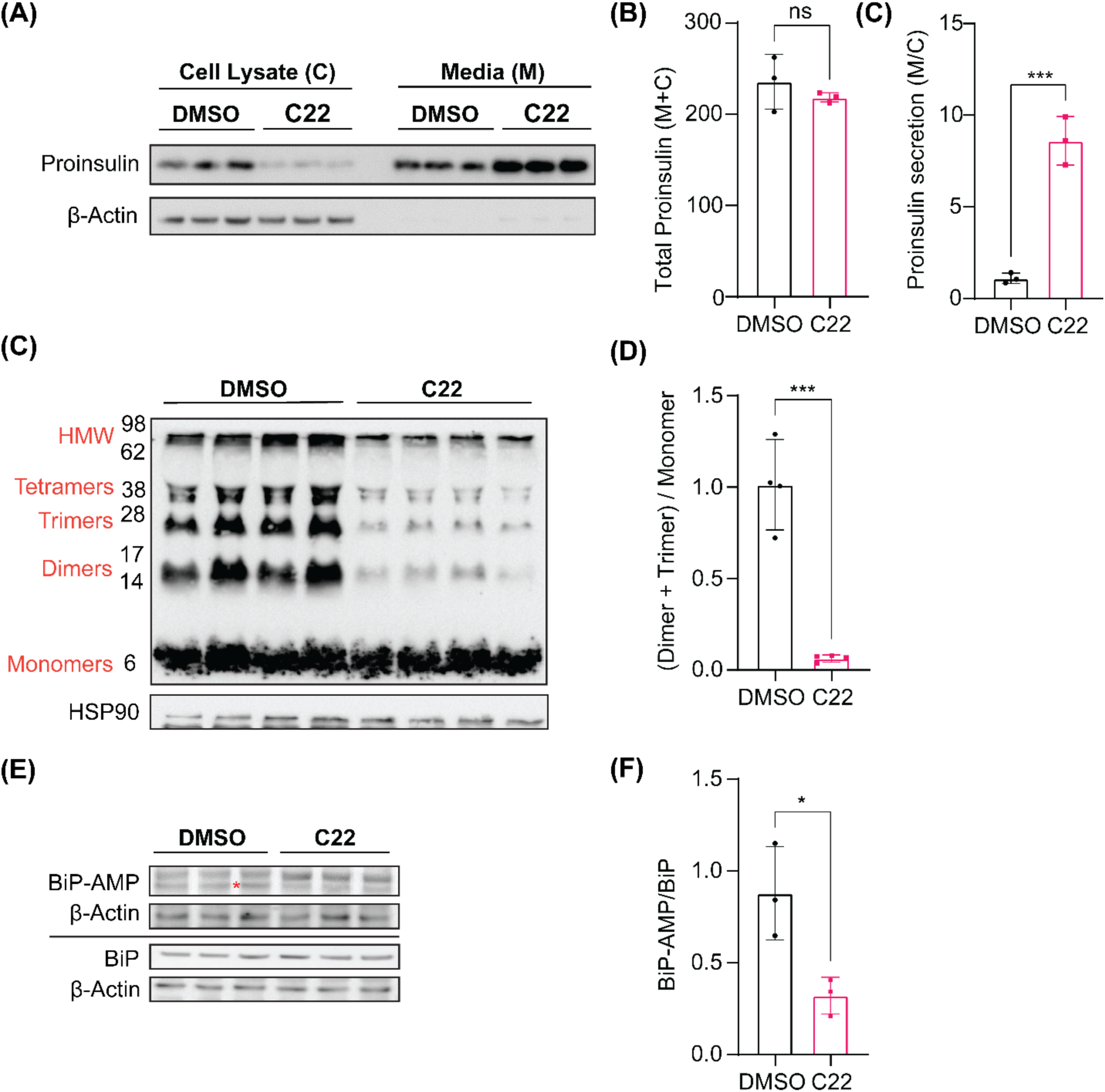
Effect of C22 and C73 on proinsulin secretion and folding in Min6 pancreatic β-cells. (**A**) Min6 cells were treated with 20 µM C22 or DMSO for 16 hours and the media (M) and cell lysate (C) were probed for proinsulin levels. β-actin was used as the protein loading control. (**B**) Total proinsulin levels in M+C as quantified from (**A**). (**C**) Proinsulin levels in M compared to C (M/C ratio) as quantified from (**A**). (**D**) Non-reducing SDS-PAGE showing proinsulin monomer and higher order oligomers. Numbers on the left represent the protein ladder molecular weights. HSP90 was used as the protein loading control. (**E**) Quantification of (**D**). (**F**) Min6 cells treated with C22 or DMSO were lysed and probed for AMPylated BiP levels. β-actin was used as the protein loading control. * indicates the protein band used for quantification purposes; the upper protein band is non-specific for BiP AMPylation (**G**) Quantification of (**F**). Statistical significance between control and treated groups was assessed by performing an unpaired t-test with Welch’s correction. Data are presented as mean ± S.D.

## Discussion

In this study, we define a pair of small molecules, C22 and C73, as novel FICD inhibitors with limited cytotoxicity. We show that these molecules inhibit BiP AMPylation by both wild-type and pathogenic FICD variants and highlight their potential to improve pro-insulin folding and secretion in pancreatic β-cells. We further demonstrate that the compounds are amendable to rational medicinal chemistry-based improvements and present a proposed mode of target engagement supported by *in-silico* and cell-based work.

Recent advances in our understanding of FICD biology provide a compelling premise for the development of FICD inhibitors for therapeutic considerations. FICD-mediated cycles of BiP AMPylation and deAMPylation regulate ER homeostasis. Pathogenic mutations in the FICD active site disrupt this equilibrium, leading to loss of deAMPylation function, which results in the accumulation of AMPylated BiP^21,22^. Additional work indicates that loss of endogenous FICD activity mitigates pressure overload-induced cardiac hypertrophy by inducing a robust UPR^ER^ response and enhancing ER-phagy in cardiomyocytes^25^. In this study, we show direct evidence for enhanced proinsulin processing in response to FICD inhibition. In all these scenarios, FICD inhibitors are expected to provide immediate benefit by reducing BiP AMPylation to restore and/or boost UPR^ER^ signaling and ER homeostasis. A small number of putative *in vitro* FICD inhibitors have previously been described^54–56^. Unlike compounds C22, and its sodium salt, C73, however, these molecules were not tested against endogenous or pathogenic FICD variants.

C22 and C73 are thus the first tool compounds to efficiently inhibit FICD in tissue culture models. C22, a halogenated salicylanilide, further shows functional group flexibility in the biaryl region of the scaffold which amends itself for rational improvement.

Using a combination of *in-silico* MD simulations and cell-based assays, we provide evidence that FICD inhibitors C22 and 73 stably bind dimeric FICD, preferentially at the smaller dimeric interface or the TPR-II domain. We propose that this interaction prevents BiP AMPylation by either blocking the switch to an AMPylation-competent, monomeric state or abrogating BiP binding to FICD. The compounds also bind monomeric FICD, which inhibits BiP AMPylation (**Figure 7**). The finding that C22 and C73 moderately inhibit FICD-mediated BiP deAMPylation further suggests that the compounds may abrogate the transition from an AMPylation-competent monomer to a deAMPylation-competent dimeric state. Determining the structure of C22-bound FICD will be a critical next step to confirm this proposed mode of action.

**Figure 7:**
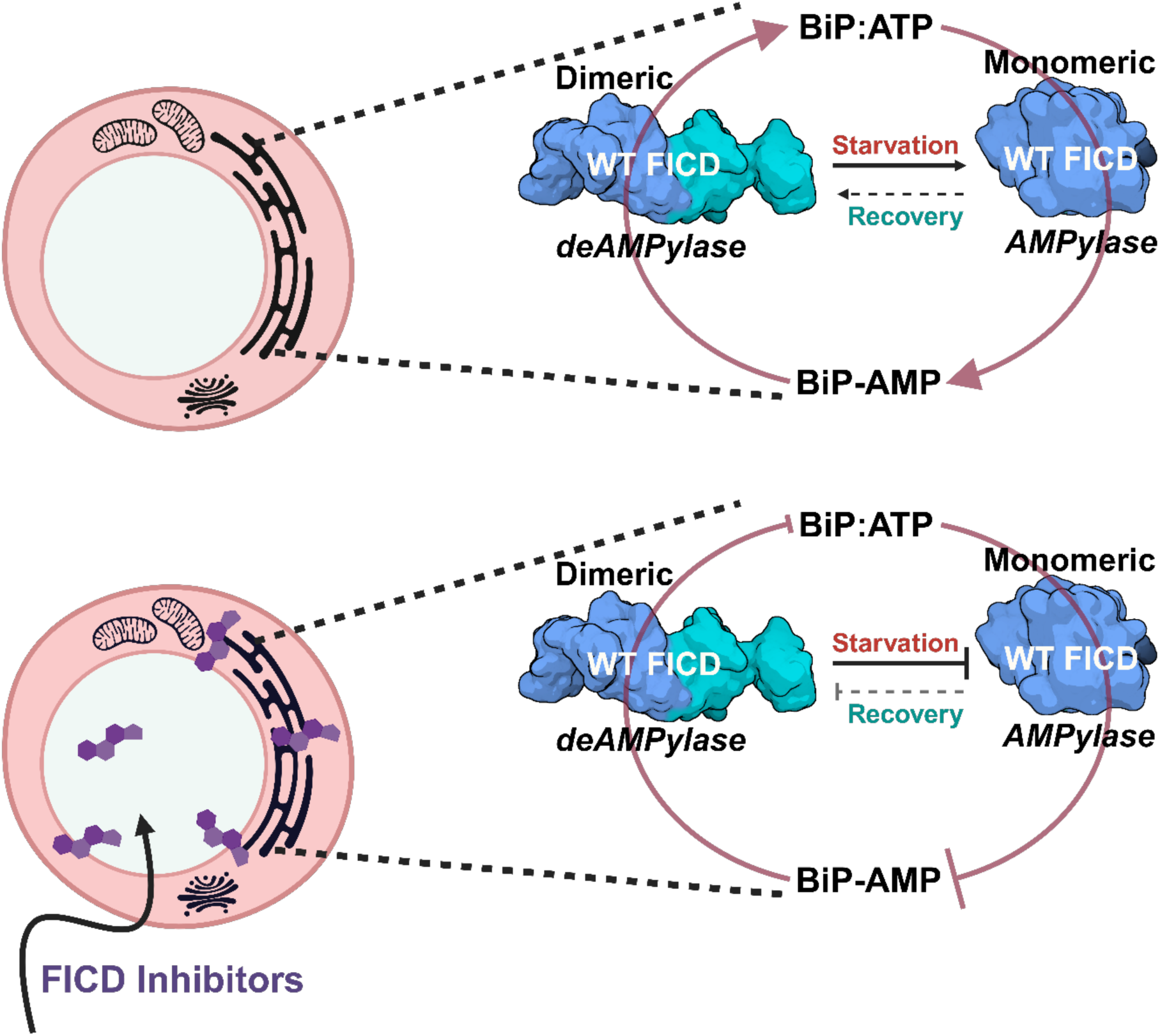
Schematic representation of the effect of FICD inhibitors on bifunctional WT FICD enzyme activity in A549 cells. PBS induced starvation triggers the reversible transition of dimeric, deAMPylation-competent FICD to a monomeric, AMPylation-competent state, resulting in the accumulation of non-functional, AMPylated BiP. When PBS is removed, the AMPylation-competent FICD reverts to its deAMPylase conformation, thereby increasing the pool of deAMPylated, active BiP (ATP-bound). FICD inhibitors strongly suppress the transition of deAMPylation-competent to AMPylation-competent states while weakly inhibiting the opposite. Strong and weak suppression of FICD function is represented by the size of the blunt arrows. Dimeric and monomeric FICD are cartoon representations of their PDB structures 4U04 and 6I7J, respectively. The schematic was made using BioRender.

Using pancreatic cells, we show that FICD inhibitor C22 enhances anterograde trafficking and proinsulin folding while reducing aggregated or misfolded proinsulin. These results are significant, as to the best of our knowledge, no other small molecule with similar capacity to improve proinsulin processing is described. Future proof-of-concept studies using in vivo models for proinsulin misfolding will provide first evidence for the therapeutic potential of using FICD inhibitors to mitigate autosomal dominant diabetes.

### Limitations of the study

This study takes the first steps towards understanding the structure-activity relationship (SAR) between FICD and compounds C22 and C73. The described SAR-based analog testing represents a glance at the potential of SAR-based medical chemistry to further improve compound efficacy. Additional efforts will likely lead to the development of a more potent lead compound. We also failed to source compound C34 in reasonable quantities preventing the testing of this most promising molecule in more assays. Testing the compounds in more cell lines, stress conditions, and ultimately, *in vivo* are critical next steps towards defining their potential for future clinical use.

## Significance

This study identifies two cell-permeable FICD inhibitors, C22 and C73, which inhibit FICD-mediated BiP AMPylation while exhibiting low cytotoxicity. Both compounds inhibit wild-type and pathologic FICD variants. This is significant, considering that the number of identified pathologic FICD variants is increasing but small molecules to target these mutant enzymes are lacking.

Our *in silico* docking work provides mechanistic insights into the mode of action of C22 and C73, confirming the proposed model that FICD oligomerization is critical to controlling its AMPylation/deAMPylation activity. Our study also demonstrates that targeting FICD improves proinsulin folding and secretion. These results establish a first link between FICD activity and proinsulin processing in the ER, and highligh a promising new application for FICD inhibitors to improve proinsulin processing in β cells. Taken as a whole, our study confirms FICD as a druggable enzyme and provides critical support for considering FICD as a target for multiple clinical indications.

## Acknowledgments

We thank the members of the Truttmann lab for helpful comments and discussion. William Giblin is acknowledged for proof-reading the manuscript draft. We are grateful for support by the Center for Chemical Genomics (CCG) at the University of Michigan Life Sciences Institute. MCT is supported by grant 1R35GM142561. CLB acknowledges funding from the NIH through R35GM130597. This work was partially supported by the Pandemic Research Recovery grant awarded to BKC by the Medical School Office of Research at the University of Michigan.

## Author Contributions

MCT supervised the project. BKC, AC, MA, JR, CLB and PDA designed and planned the experiments. BKC, AC, MA and SML conducted the experiments. BKC and MCT wrote the manuscript and all authors edited and approved the final manuscript.

