## Supplementary Figures and Tables for "Small molecule FICD inhibitors suppress endogenous and pathologic FICD-mediated protein AMPylation"

| Name | Sequence |
|---|---|
| *ficd F* | TGAAGCCATTGGTATGAGCA |
| *ficd R* | GCGGTTGGGGTTCTGAC |
| *xbp1(total) F* | GGCATCCTGGCTTGCCTCCA |
| *xbp1 (total) R* | GCCCCCTCAGCAGGTGTTCC |
| *actb F* | ACAGAGCCTCGCCTTTGCC |
| *actb R* | GATATCATCATCCATGGTGAGCTGG |
| *bip F* | GGATCATCAACGAGCCTACGG |
| *bip R* | AGACACATCGAAGGTTCCGC |
| *xbp1 (spliced) F* | CCGCAGCAGGTGCAGG |
| *xbp1 (spliced) R* | GAGTCAATACCGCCAGAATCCA |
| *atf4 F* | GTTCTCCAGCGACAAGGCTA |
| *atf4 R* | GAGAAGGCATCCTCCTTGCT |
| *casp4 F* | TTCCCTATGGCAGAAGGCAAC |
| *casp4 R* | GCCATGACCCGAACTTTGTC |
| *casp6 F* | GAGGAGGGCAAGGTGTCTG |
| *casp6 R* | TGTTTTCTTCCCCACCTGC |

| | |
|---|---|
| *atf6α F* | GCTCTGGAACAGGGCTCAAA |
| *atf6α R* | CTCCCTGAGTTCCTGCTGAT |

**Supplementary Table S1. List of qPCR primers used in the study**. Primer pairs were obtained from IDT, and they are listed as 5' to 3'. F and R refer to forward and reverse primer sequences, respectively. The primers were designed using the National Centre for Biotechnology Information (NCBI) "pick primer" tool and validated for specificity and amplicon size using an in-silico PCR webtool from UCSC genome browser (https://genome.ucsc.edu/cgi-bin/hgPcr).

| # | Compound Name | $IC_{50}$ (μM) | 95 % CI (profile likelihood) |
|---|---|---|---|
| 1 | C47 | 18.3 | 13.9 - 24.2 |
| 2 | C22 | 7.6 | 5.9 - 9.7 |
| 3 | C73 | 8.2 | 6.3 - 10.8 |
| 4 | C522 | 6.1 | 4.6 - 8.2 |
| 5 | C83 | 14.6 | 12.2 - 17.3 |
| 6 | C84 | 4.49 | 3.0 - 6.7 |
| 7 | C53 | 5.7 | 5 – 6.5 |

**Supplementary Table S2: $IC_{50}$ values of top 7 compounds.** 1 μM $FICD^{E234G}$ was incubated with varying concentrations of the top 7 putative FICD inhibitors for 10-15 mins at RT, following which FL-ATP was added. The reaction was incubated for 90 mins at 37°C in the dark. $IC_{50}$ values (μM), as shown in the Supplementary Table, were obtained by fitting the dose response values using a non-linear regression method in GraphPad Prism. CI: Confidence Interval.

| # | Cell type | $LD_{50}$ (μM) | 95 % CI (profile likelihood) |
|---|---|---|---|
| 1 | HeLa | 133 (C22) | 116 to 153 |
| 2 | | 148 (C73) | 128 to 172 |
| 3 | SK-N-SH | 92.8 (C22) | 78.1 to 111 |
| 4 | | 98.8 (C73) | 76.3 to 134 |
| 5 | A549 | 190.3 (C22) | 166.0 to 227.4 |
| 6 | | 167.1 (C73) | 115.7 to 320.9 |
| 7 | Neonatal Murine cardiomyocytes | 88.6 (C22) | N/A |
| 8 | | 95.1 (C73) | N/A |

**Supplementary Table S3: $LD_{50}$ values of C22 and C73 in human cells.** Various human cells were incubated with compounds C22 and C73 for 24 hours and cell viability was assessed using the MTT assay. $LD_{50}$ values (μM), as shown in the Supplementary Table, were obtained by fitting the dose response values using a non-linear regression method in GraphPad Prism. N/A indicates that Prism was unable to calculate confidence interval values (CI) for the non-linear regression fit and hence the $LD_{50}$ value should be interpreted with caution.

| FICD variant | Compound | Rank | Contacts (< 5Å) | | FACTS-score ( mean +/- S.D.) |
|---|---|---|---|---|---|
| | | | *Amino Acid* | *Residue* | |
| WT dimer | C22 | 1 | SER | 288 | -28.8 6 +/-4.89 |
| | | | LEU | 291 | |
| | | | GLU | 292 | |
| | | | ARG | 295 | |
| | | | ALA | 306 | |
| | | | GLY | 307 | |
| | | | SER | 288 | |
| | | | LEU | 291 | |
| | | | GLU | 292 | |
| | | | ARG | 295 | |
| | | | ALA | 306 | |
| | | | GLY | 307 | |
| | | 2 | THR | 164 | -20.83 +/ 2.75 |
| | | | THR | 168 | |
| | | | PRO | 171 | |
| | | | TYR | 172 | |
| | | | ARG | 180 | |
| | | | GLY | 409 | |
| | | | ASP | 410 | |
| WT dimer | C73 | 1 | LEU | 167 | -17.9 +/- 5.25 |
| | | | PRO | 171 | |
| | | | ARG | 180 | |
| | | | GLY | 409 | |
| | | | ASP | 410 | |
| | | | ARG | 412 | |
| | | 2 | LEU | 291 | -15.23 +/- 9.99 |
| | | | GLU | 292 | |
| | | | ARG | 295 | |
| | | | ALA | 306 | |
| | | | GLY | 307 | |
| | | | ARG | 308 | |
| | | | LEU | 291 | |
| | | | ARG | 295 | |
| | | | ALA | 306 | |
| | | | GLY | 307 | |
| | | | PHE | 309 | |

**Supplementary Table S4: Binding sites and docking scores of FICD inhibitors bound to dimeric WT FICD.** Interacting amino acid residues for the top 2 binding poses are listed along with their respective FACTS docking score. All interacting residues are within a 5Å radii of the compound(s) binding cavity.

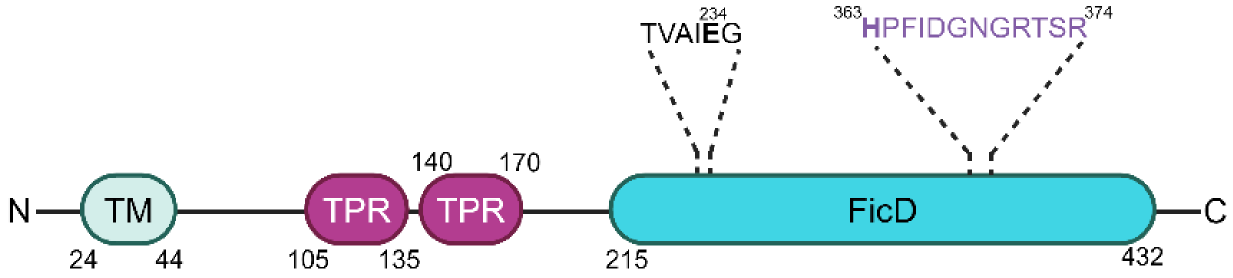


**Supplementary Figure S1: Domain organization of the human AMPylase FICD**. Schematic depiction of FICD protein domain architecture showing transmembrane domain (TM), tetratricopeptide domain (TPR) and Fic domain (FicD). The glutamate-containing inhibitory motif and histidine-containing Fic motif are depicted in black and purple, respectively.

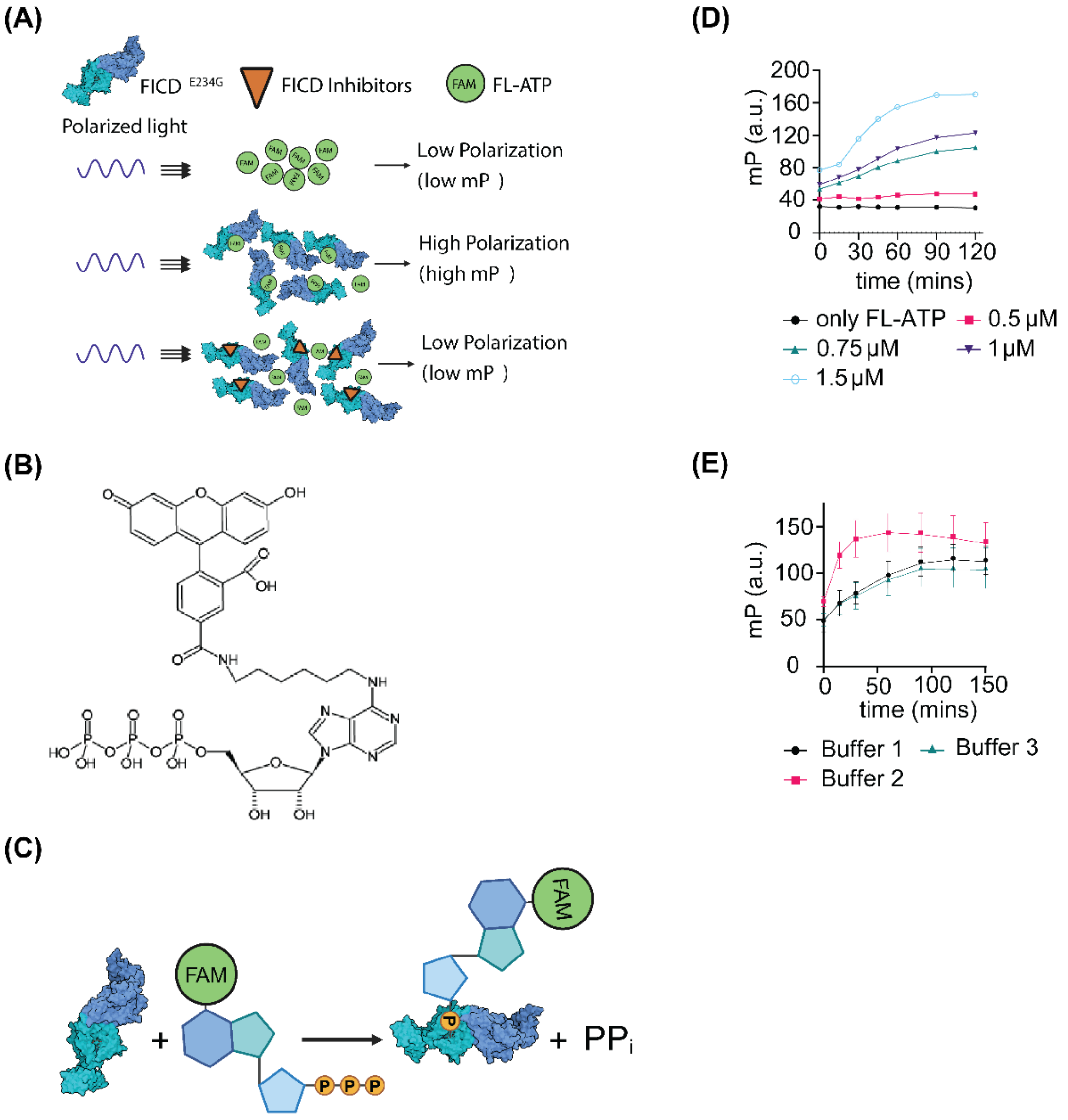


**Supplementary Figure S2**. **FP assay optimization**. (**A**) FP assay schematic. Binding of FL-ATP to FICD$^{E234G}$ reduces the degree of freedom of the fluorophore and slows the rotation of the protein-fluorophore complex, resulting in a significant increase in the emission of polarized light, i.e. FP. FICD$^{E234G}$ auto-AMPylation Increases FP; putative FICD inhibitors are expected to decrease FP. (**B**) Structure of FL-ATP (N6-(6-Aminohexyl)-ATP-5-FAM). (**C**) FICD$^{E234G}$ binds to FL-ATP and catalyzes auto-AMPylation. (**D**) FICD$^{E234G}$ at indicated concentrations was incubated with 250 nM FL-ATP. FP was measured at indicated timepoints. Wells containing only FL-ATP in AMPylation buffer served as negative controls. Quadruplicate measurements were averaged for each FICD concentration and plotted as a function of time. (**E**) 1 μM FICD$^{E234G}$ was incubated with 250 nM FL-ATP in buffer 1 (50 mM HEPES pH 7.5, 150 mM NaCl, 10 mM $MgCl_2$, 1 mM DTT and 1 mM EDTA) or buffer 2 (50 mM HEPES pH 7.5, 150 mM NaCl, 10 mM $MnCl_2$, 1 mM DTT and 1 mM EDTA) or buffer 3 (50 mM HEPES pH 7.5, 150 mM NaCl, 10 mM $MgCl_2$ and 1 mM DTT. Wells containing only FL-ATP in AMPylation buffer served as negative controls. Quadruplicate measurements were averaged for each time point and buffer. Error bars represent standard deviation (S.D.).

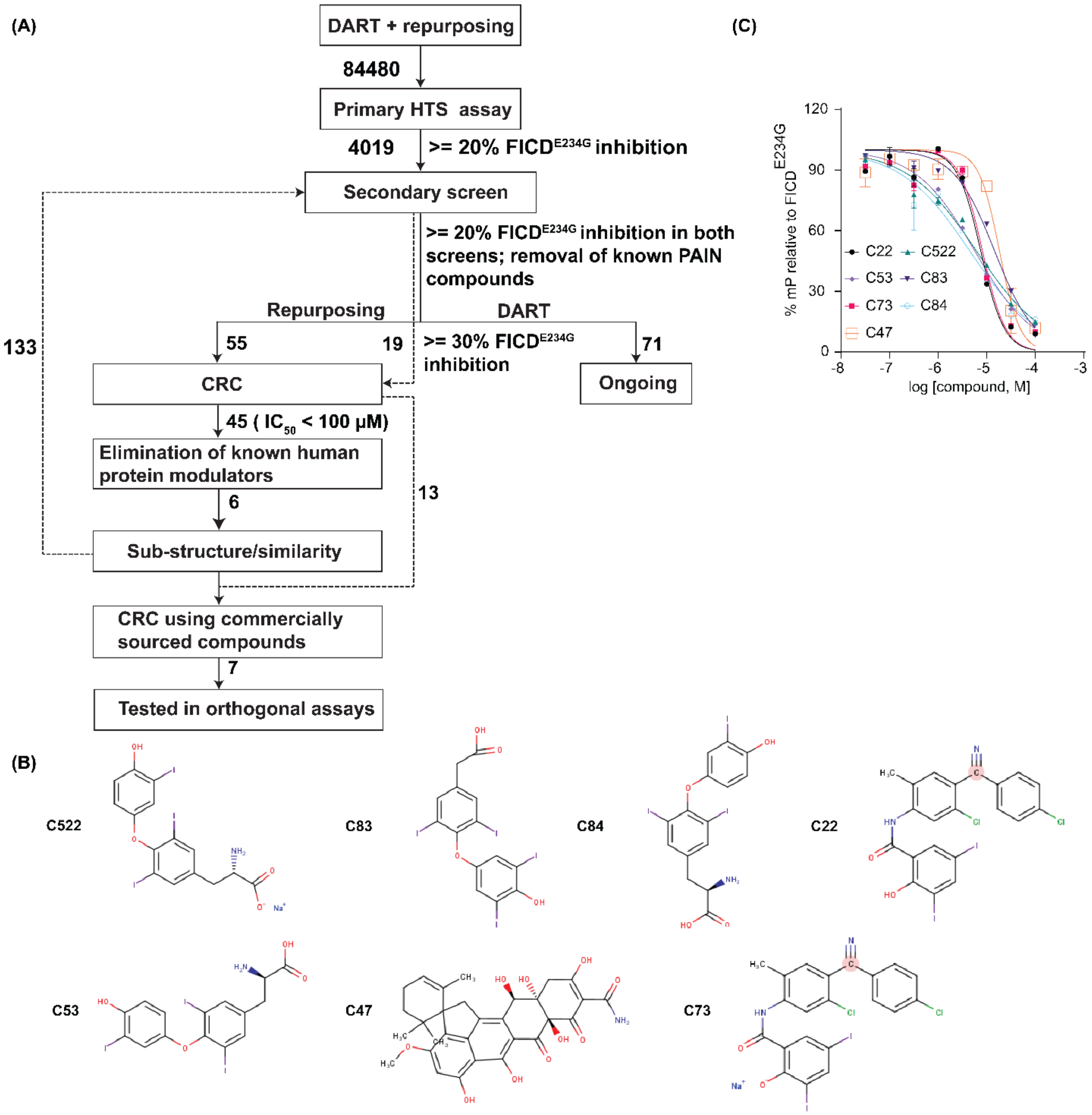


**Supplementary Figure S3: Concentration response assay for FICD inhibitors**. (**A**) Drug discovery and development pipeline. Numbers in italics represent the number of compounds selected for the next step in the pipeline. Only compounds that exhibited a sigmoidal or linear concentration dependence and showed an $IC_{50}$ < 100 μM were selected for the next steps. Sub-structure search was carried out using a 40% structural similarity cut-off. (**B**) Chemical structures of the top 7 lead compounds drawn using MarvinSketch (v 24.1.2) (**C**) Concentration Response assay for the top 7 compounds. $IC_{50}$ values were obtained by fitting the data using a non-linear regression method in GraphPad Prism.

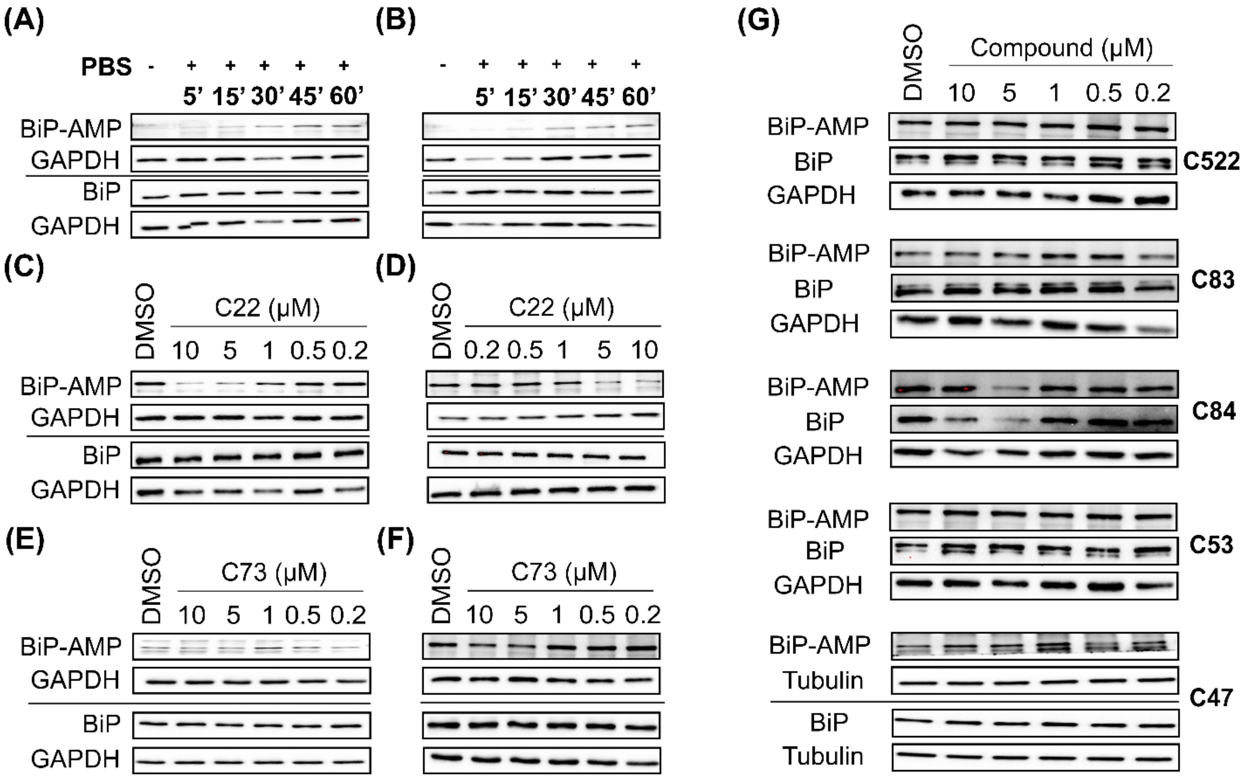


**Supplementary Figure S4: Evaluating concentration response of FICD inhibitors in PBS treated A549 cells.** (**A-B**) AMPylated and total BiP levels of A549 cells maintained in GM or treated with sterile PBS for the indicated time points. (**C-D**) AMPylated and total BiP levels of A549 cells treated with PBS for one hour in the presence of either DMSO (control) or C22 (at indicated concentrations). (**E-F**) AMPylated and total BiP levels of A549 cells treated with PBS for one hour in the presence of either DMSO (control) or C73 (at indicated concentrations). In all experiments, equal amounts of protein were loaded onto two separate reducing SDS-PAGE gels and protein transferred onto PVDF membranes. One was probed for AMPylated BiP while the other for total BiP. GAPDH was used as the protein loading control. (**G**) AMPylated and total BiP levels of A549 cells exposed to PBS supplemented with either 0.5% (v/v) DMSO or compounds C47, C522, C83, C84 and C53 for one hour. GAPDH or α-Tubulin was used as the protein loading control.

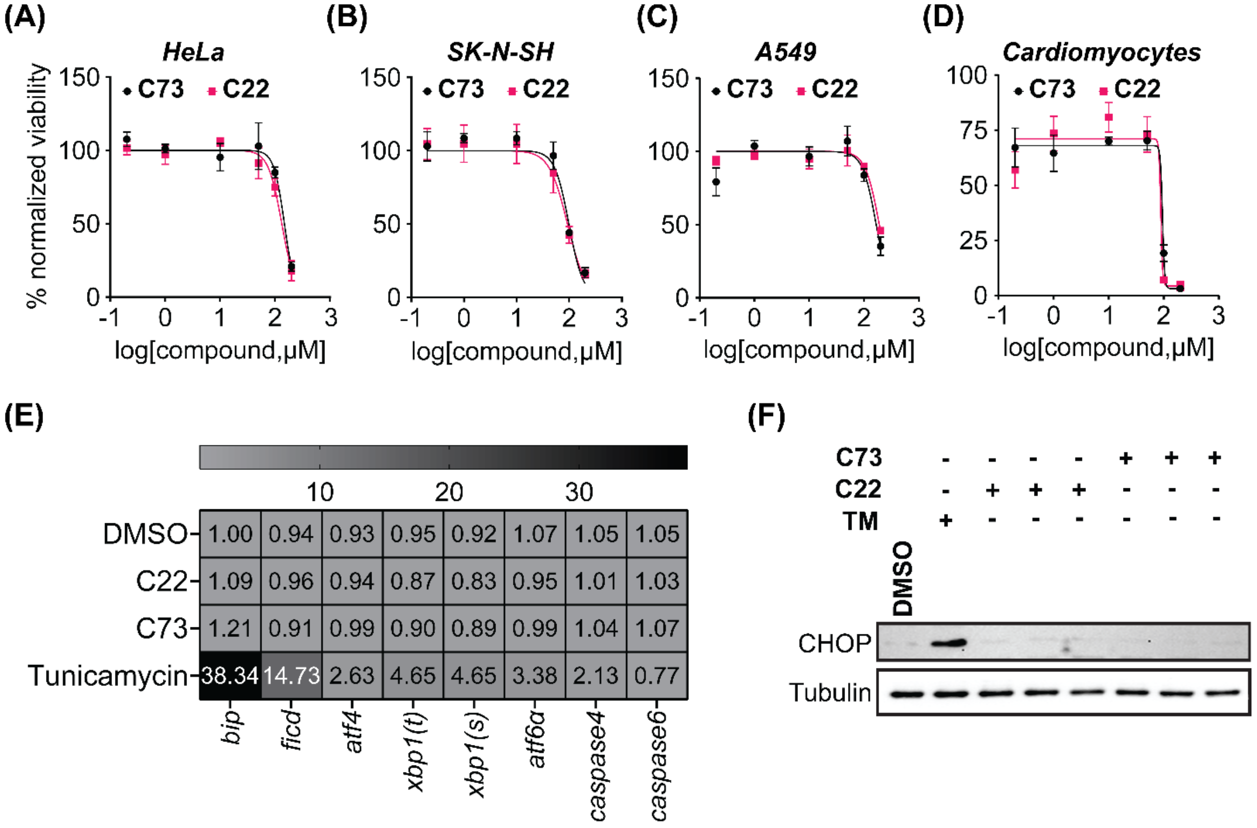


**Supplementary Figure S5: Evaluation of C22 and C73 induced cytotoxicity and UPR$^{ER}$ in human cells.** MTT assay was carried out in (**A**) HeLa, (**B**) SK-N-SH, (**C**) A549 cell lines, and (**D**) neonatal C57BL/6J cardiomyocytes to evaluate compound cytotoxicity. 0.2, 1, 10, 50, 100 and 200 μM FICD inhibitors were used for all cytotoxicity experiments. Each dot represents the mean of three independent biological replicas, with each biological replica consisting of three technical replicas. The arrows represent S.D. of mean values (**E**) A549 cells were incubated with 5 μM C22 or C73 for 24 hours in GM. Cells exposed to an equivalent amount of DMSO or 2 μg/ml of tunicamycin for 24 hours acted as the negative and positive controls, respectively. Ribonucleic acid (RNA) extracted from cell lysates were reverse transcribed to obtain complementary DNA (cDNA) and 20ug cDNA was used for a single qPCR reaction. Fold changes in gene expression are denoted by the numbers inside each cell of the heat map. The numbers represent the mean of three independent biological replicas, with each biological replica consisting of three technical replicas. Gene expression levels were normalized to *ACTB* which was used as the housekeeping control. (**F**) A549 cells were incubated with 5 μM C22 or

C73, or an equivalent amount of DMSO for 24 hours in GM. Cells exposed to 25 µg/ml tunicamycin (TM) for 2 hours acted as a positive control. Cell lysates, obtained from three independent biological replicas for each compound, were probed for CHOP levels and α-tubulin was used as the loading control.

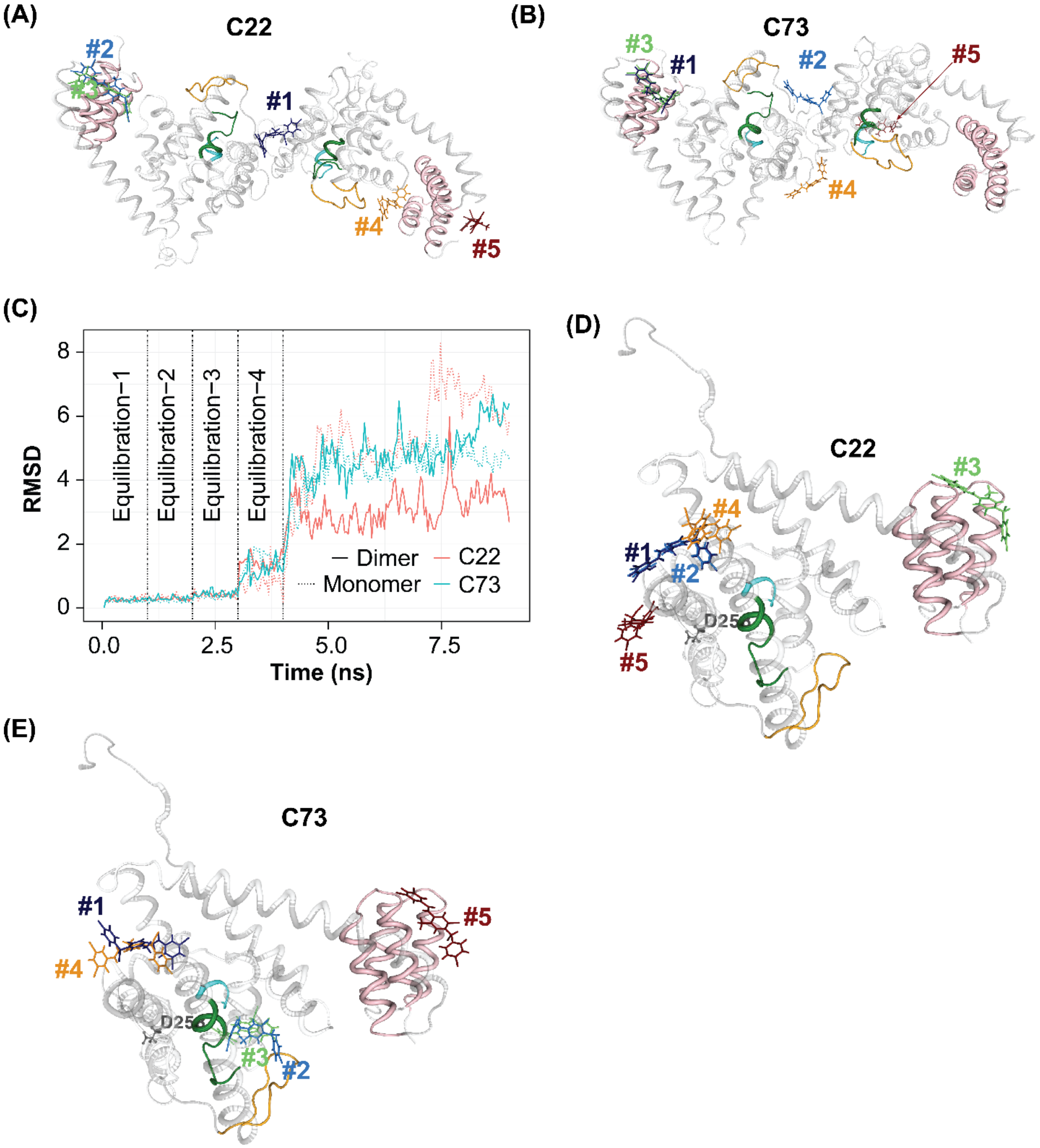

**Supplementary Figure S6:** ***In-silico*** **docking of FICD inhibitors to dimeric and monomeric FICD**. (**A-B**) The top 5 binding sites within the apo dimeric WT FICD (PDB ID: 4U04) for C22 (**A**) and C73 (**B**). (**C**) Structural sidechain RMSD of the compounds bound to FICD variants during the 10 ns MD simulations. Only the heavy atom RMSD is computed, and the individual snapshots are aligned to minimize the translational and rotational deviation of the receptor's/enzyme's backbone. (**D-E**) The top 5 binding sites within the monomeric $FICD^{L258D}$ (PDB ID: 6I7J) for C22 (**D**) and C73 (**E**). TPR domains are highlighted in pink. The catalytic flap and the inhibitory α- helix are highlighted in yellow and green, respectively.

(A)

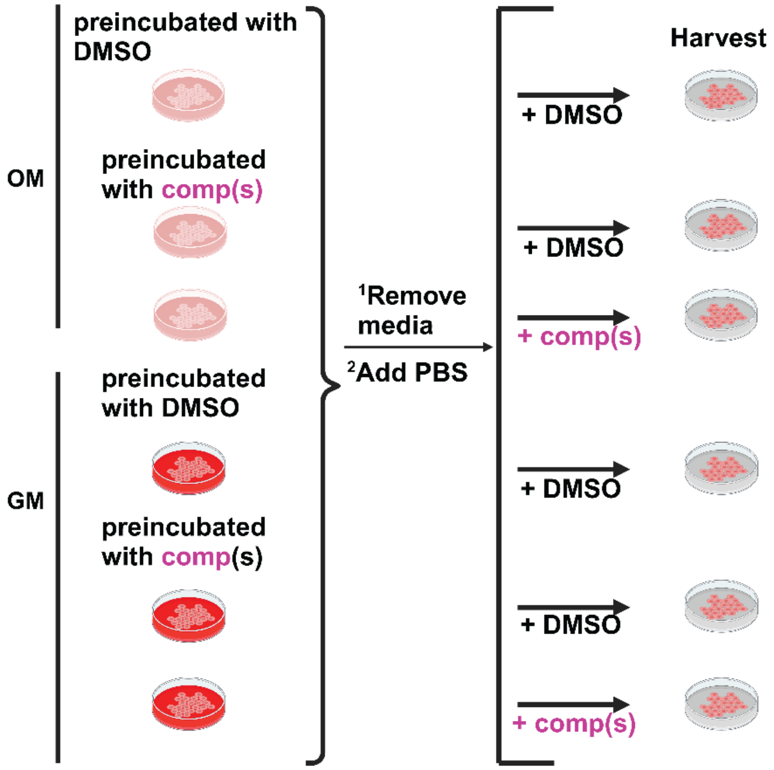


(B)

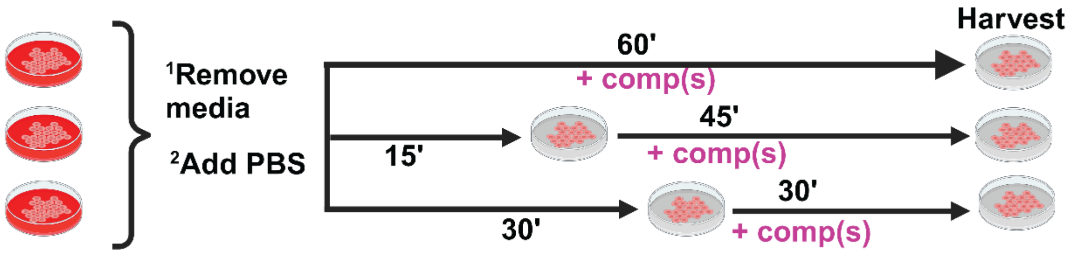


**Supplementary Figure S7: Schematics depicting cell treatment regime with various media.** (**A**) Schematic depicting treatment regime of A549 cells preincubated with the compounds in either Opti-MEM (OM) or GM and subsequently exposed to PBS supplemented with either FICD inhibitors or DMSO. (**B**) Schematic depicting PBS treatment regime of A549 cells in the presence of FICD Inhibitors or DMSO added at the indicated time points.

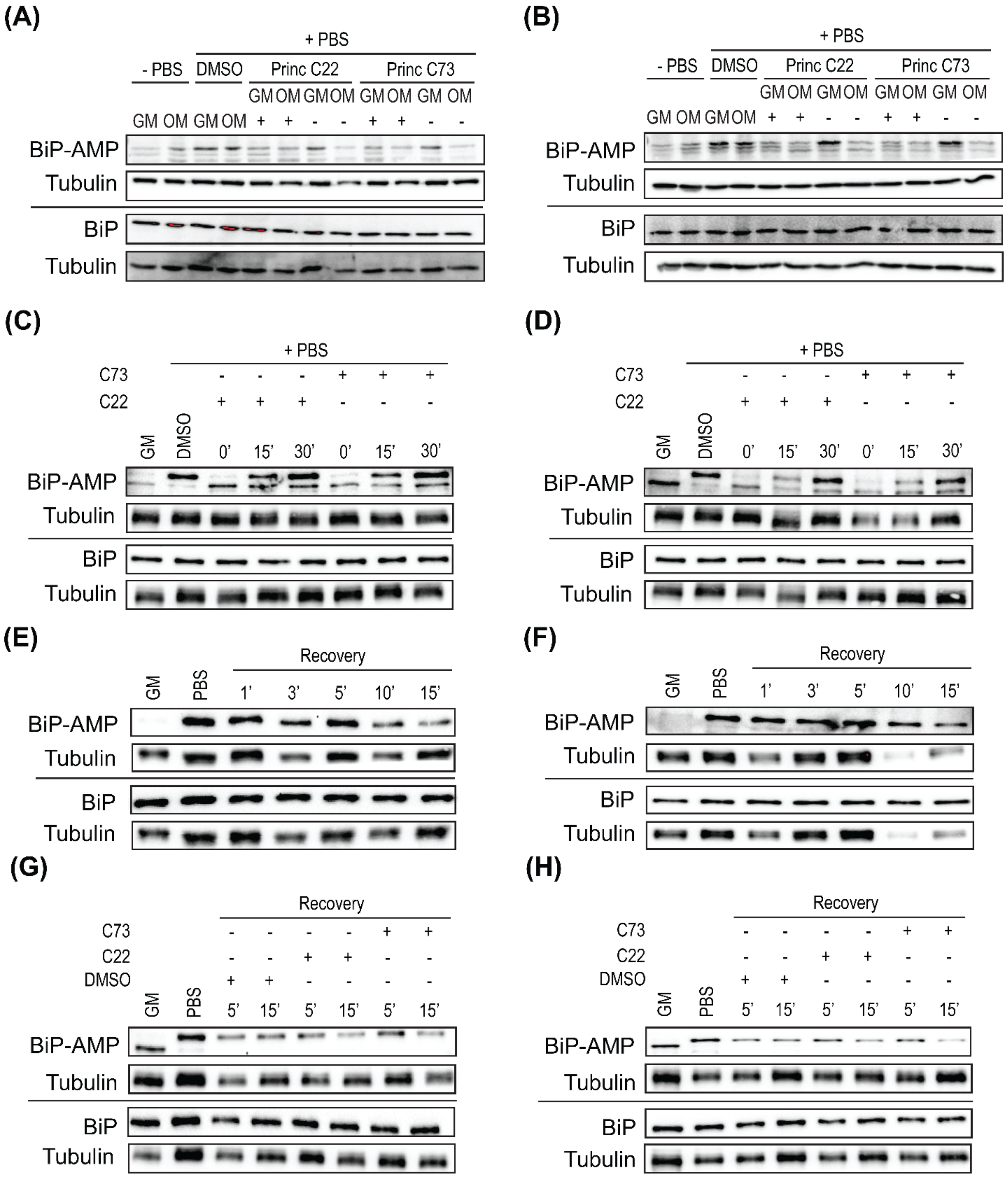


**Supplementary Figure S8: The effect of FICD inhibitors on deAMPylation and AMPylation competent FICD states**. (**A-B**) AMPylated and total BiP levels in A549 cells preincubated with the 10 µM FICD inhibitors in GM or OM and subsequently exposed to PBS supplemented with 10 µM FICD inhibitors for 15 mins. Cells preincubated with 0.5% (v/v) DMSO and exposed to

PBS supplemented with 0.5% (v/v) DMSO served as positive controls while cells grown in GM or OM supplemented with 0.5% (v/v) DMSO but not exposed to PBS served as negative controls. From left: Lanes 1-2 represent negative controls, lanes 3-4 represent positive controls, lanes 5-6 represent cells preincubated with C22 and exposed to PBS supplemented with C22 while lanes 7-8 represent cells preincubated with but not subsequently exposed to C22 during PBS treatment. Lanes 8-12 follow the exact order as lanes 5-8 but represent cells treated with C73. (**C-D**) AMPylated and total BiP levels cells exposed to PBS supplemented with 10 μM C22 or C73 for the indicated time points. 0', 15' and 30' represent the time at which the compounds were added to PBS. (**E-F)** AMPylated and total BiP levels of PBS treated cells incubated in GM for the indicated time points. (**G-H**) AMPylated and total BiP levels of PBS treated cells incubated in GM with or without 10 μM C22 or C73, for the indicated time points. α-Tubulin was used as the protein loading control. Schematics shown in (**A**) and (**B**) were made using BioRender.

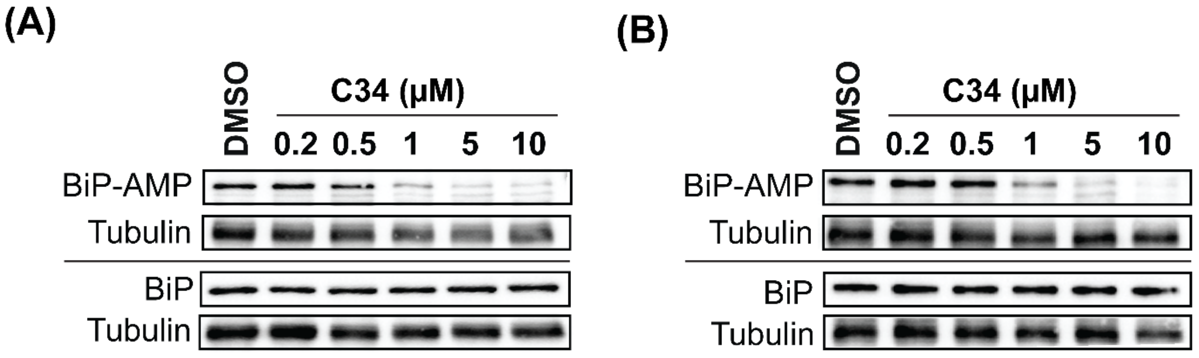


**Supplementary Figure S9: Dose response assessment of closantel analog C34**. (**A-B**) AMPylated and total BiP levels of A549 cells treated with PBS for one hour in the presence of either 0.5% (v/v) DMSO (control) or compound C34 (at indicated concentrations). α-Tubulin was used as the protein loading control.

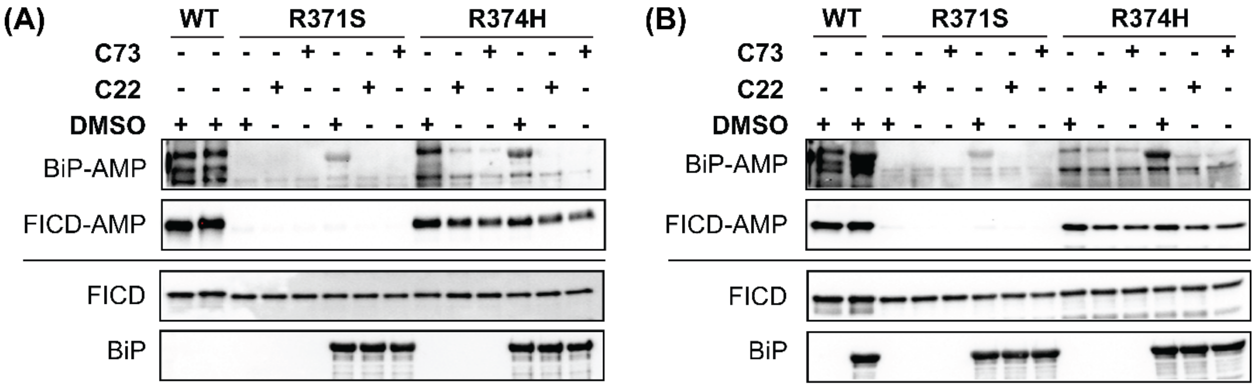


**Supplementary Figure S10: Effect of C22 and C73 on pathologic FICD-mediated BiP AMPylation *in-vitro***. (**A-B**) AMPylated and total levels of recombinant human BiP in the presence of FICD inhibitors or DMSO. WT FICD (+/-) BiP served as negative controls.

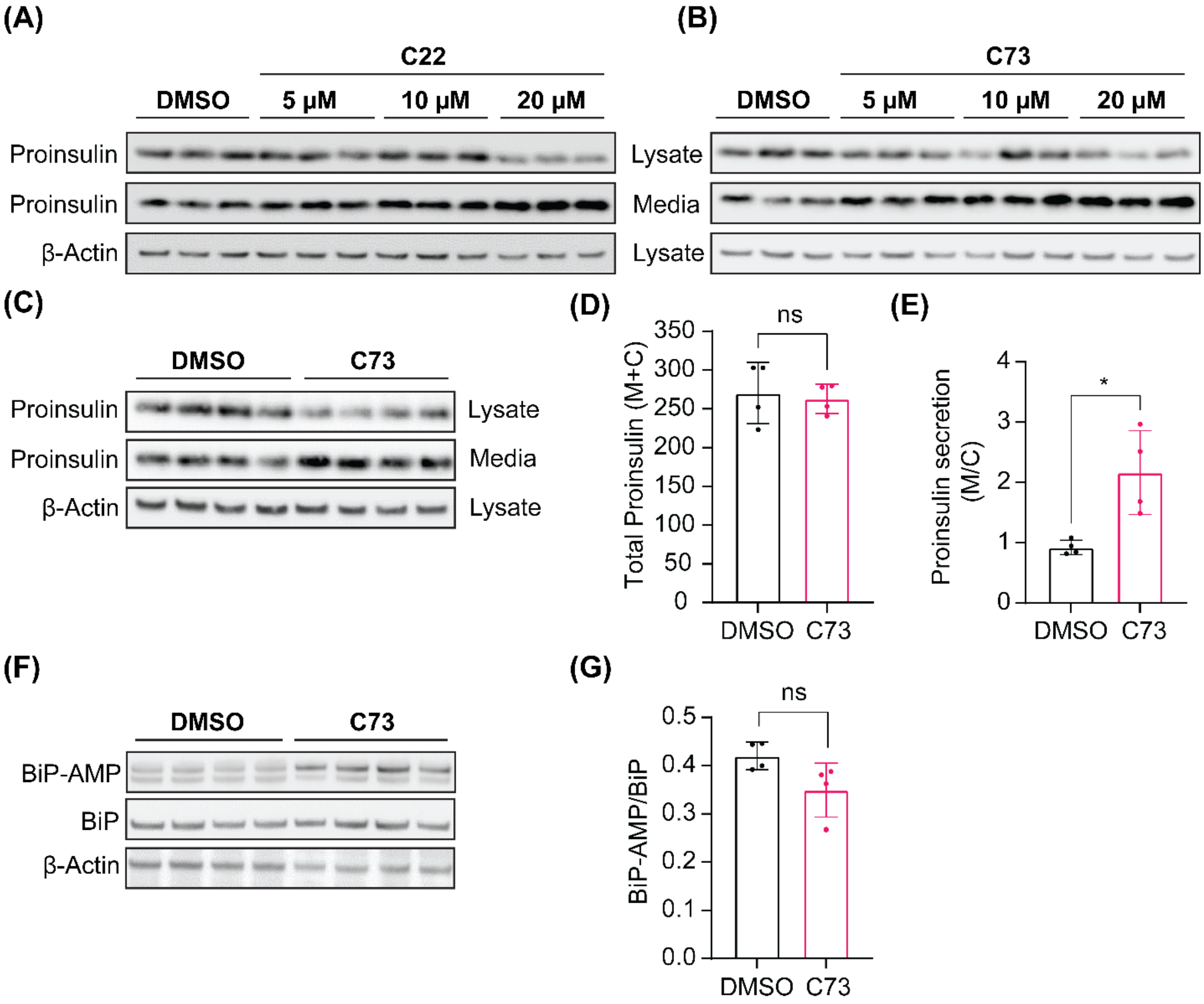

**Supplementary Figure S11: Effect of C22 and C73 on proinsulin secretion and folding in Min6 pancreatic β-cells**. (**A-B**) Cells were exposed to indicated concentrations of C22 (**A**) and C73 (**B**) or DMSO for 16 hours and the media (M) and cell lysate (C) were probed for proinsulin levels. (**C**) Min6 cells were incubated with 20 μM C73 or DMSO for 16 hours and probed for proinsulin levels. (**D**) Total proinsulin levels in M+C as quantified from (**C**). (**E**) Proinsulin levels in M compared to C (M/C ratio) as quantified from (**C**). (**F**) Min6 cells were treated with 20 μM C73 or DMSO for 16 hours and probed for AMPylated BiP levels. (**G**) Quantification of (**F**). β-actin was used as the protein loading control in all experiments. Statistical significance between control and treated groups was assessed by performing an unpaired t-test with Welch's correction. Data are presented as mean ± S.D.
